# Microvesicles Transfer Mitochondria and Increase Mitochondrial Function in Brain Endothelial Cells

**DOI:** 10.1101/2021.04.10.439214

**Authors:** Anisha D’Souza, Amelia Burch, Kandarp M. Dave, Aravind Sreeram, Michael J. Reynolds, Duncan X. Dobbins, Yashika S. Kamte, Wanzhu Zhao, Courtney Sabatelle, Gina M. Joy, Vishal Soman, Uma R. Chandran, Sruti S. Shiva, Nidia Quillinan, Paco S. Herson, Devika S Manickam

## Abstract

We have demonstrated, for the first time that microvesicles, a sub-type of extracellular vesicles (EVs) derived from hCMEC/D3: a human brain endothelial cell (BEC) line transfer polarized mitochondria to recipient BECs in culture and to neurons in mice acute brain cortical and hippocampal slices. This mitochondrial transfer increased ATP levels by 100 to 200-fold (relative to untreated cells) in the recipient BECs exposed to oxygen-glucose deprivation, an *in vitro* model of cerebral ischemia. We have also demonstrated that transfer of microvesicles, the larger EV fraction, but not exosomes resulted in increased mitochondrial function in hypoxic endothelial cultures. Gene ontology and pathway enrichment analysis of EVs revealed a very high association to glycolysis-related processes. In comparison to heterotypic macrophage-derived EVs, BEC-derived EVs demonstrated a greater selectivity to transfer mitochondria and increase endothelial cell survival under ischemic conditions.

**Highlights:**

- Microvesicles transfer mitochondria to endothelial cells and brain slice neurons
- Mitochondrial transfer increased ATP in ischemic brain endothelial cells (BECs)
- Transfer of microvesicles increased mitochondrial function in hypoxic BECs
- Transfer of exosomes did not affect mitochondrial function in hypoxic BECs
- Homotypic BEC-derived EVs result in greater ATP levels in the recipient BECs

## 1. Introduction

The documented roles of cell-secreted extracellular vesicles (EVs) in intercellular communication via the transfer of their innate cargo make them attractive drug delivery carriers [1–11]. The subtypes of EVs vary in particle diameters, among other factors, with the larger, 100- 1000 nm microvesicles (MVs), and the smaller, 50 – 150 nm exosomes (EXOs) being secreted via different biogenesis pathways [1, 6–8, 12–14]. The lower immunogenicity of EVs, their increased stability in systemic circulation [15–19] and their ability to cross biological barriers make them attractive candidates for the delivery of biologics like nucleic acids and proteins [20–22]. Kanada *et al.* loaded plasmid DNA [23] and minicircle DNA [24] in EVs. They reported that MVs were a suitable carrier for pDNA as the EXOs failed to show the gene expression in the recipient HEK293 cells [23]. Lamichhane *et al.* also loaded DNA into EVs via electroporation and found that the loading efficiency of DNA constructs are dependent upon the size of DNA. Plasmid DNA and linear DNA greater than 1000 bp could not be loaded in EVs with a reported μg of linear 250 bp dsDNA. Nevertheless, the loading efficiency of DNA was limited at 0.2 %, with poor transfection outcomes [25].

We speculated that brain endothelial cell-derived EVs may express a natural affinity to brain endothelial cells and therefore we engineered EVs derived from hCMEC/D3, a human brain endothelial cell line, for the delivery of a model plasmid DNA. Following an ischemic insult, the inflammatory response to injury is triggered by the infiltration of peripheral immune cells across the blood-brain barrier. Yuan *et al*. [21] demonstrated that naïve EVs derived from RAW 264.7 macrophages utilize the integrin lymphocyte function-associated antigen 1, intercellular adhesion molecule 1, and the carbohydrate-binding C-type lectin receptors, to interact with hCMEC/D3 monolayers. Therefore, we also tested EVs derived from a macrophage cell line, RAW 264.7, as a delivery carrier. In our previous study, we demonstrated that macrophage-derived EVs demonstrated increased luciferase transgene expression in the recipient brain endothelial cells (BECs) compared to the homotypic EVs isolated from brain endothelial cells. More interestingly, EXOs, the smaller EV fraction, showed a greater DNA loading and transfection in the recipient BECs compared to the larger MVs [26]. The above results on EXO vs. MV-mediated DNA transfection were in direct contrast to previous observations reported by Kanada *et al*. [23] albeit their study used a different cell line (HEK293). To rule out operator-induced systematic biases in the transfection of the parent BECs using pDNA, procedures pertaining to the isolation of DNA-loaded EVs, and transfection of the recipient cells were repeated from our previous studies by an independent operator to determine the DNA delivery potential of BEC- vs. macrophage-derived EVs. We also compared BEC- and RAW-derived EVs using gene ontology and pathway enrichment analysis to understand potential differences in their composition.

The presence of a rich, innate biomolecular cargo in EVs can be exploited for additive/synergistic therapeutic effects depending on the drug cargo that is packaged in these vesicles. Specifically, EVs are reported to contain mitochondria, mitochondrial proteins, or mitochondrial DNA that can be transferred from the parent/donor to recipient cells [27]. Transfer of intact mitochondria via EVs to the recipient cells has been reported, especially during stress and injury [27] and the transferred mitochondria localize within the recipient’s mitochondrial network [28], resulting in increased cellular ATP levels [29]. Depleted oxygen and nutrient supply decrease the overall cellular energy (ATP) levels and further generate reactive oxygen species resulting in mitochondrial dysfunction, decreased cell viability and trigger apoptotic endothelial and neuronal death during ischemic stroke [20, 30].We sought to harness the innate EV mitochondrial load to increase cellular energetics in ischemic endothelial cells as a potent strategy to protect the blood-brain barrier (BBB), increase its cellular energetics and limit BBB- induced dysfunction post-stroke. We studied the effects of naïve EVs (EXOs and MVs) isolated from a brain endothelial- and macrophage cell lines on the resulting ATP levels in recipient BEC exposed to oxygen-glucose-deprived conditions, mimicking ischemic stroke-like conditions *in vitro*. We demonstrated the feasibility of EVs to deliver ATP5A, an exogenous mitochondrial protein, to increase ATP levels in ischemic endothelial cells. Further, we also investigated if EVs can transfer active and functional (polarized) mitochondria to the recipient BECs and the effects of this transfer on mitochondrial function.

We have demonstrated selective differences in the nature of EV-mediated mitochondrial transfer. Transfer of mitochondria via microvesicles, the larger EV fraction, resulted in increased mitochondrial function whereas exosome-mediated transfer did not affect mitochondrial function. We have also demonstrated, for the first time, that BEC-derived microvesicles (the larger EV fraction) transfer functional, polarized mitochondria to endothelial cells in culture and neurons in mice acute brain cortical and hippocampal slices. The presented results are of high significance as they demonstrate evidence for the potential of BEC-derived EVs to increase endothelial cell survival under ischemic conditions. Secondly, the capability of MVs to transfer polarized mitochondria to endothelial- and neuronal cells in the brain slices have important implications in the context of ischemic protection.

## 2. Experimental section

### 2.1. Materials

Reporter plasmid DNA constructs encoding the firefly luciferase gene (gWIZ-Luc/Luc- pDNA, 6732 bp) and enhanced green fluorescent protein (gWiz-GFP/GFP-pDNA, 5757 bp) were purchased from Aldevron (Fargo, ND). The stock solutions of pDNA were diluted in 10 mM Tris-HCl containing 1 mM EDTA (pH 7.4) to obtain a concentration of 1 mg/mL. The concentration and purity of the diluted solutions were confirmed by measuring A_260_/A_280_ on a NanoDrop 1000 spectrophotometer (Thermo Scientific, USA). Recombinant Human Adenosine triphosphate synthase subunit alpha (ATP5A)-Glutathione S-transferase (GST) (N-term) full-length recombinant protein was obtained from Novus Biologicals Inc. (Littleton, CO).

Lipofectamine 3000 Reagent kit was purchased from Invitrogen (Carlsbad, CA). Beetle luciferin (potassium salt), and luciferase cell culture lysis 5*x* reagent were purchased from Promega (Madison, WI). Bovine lung aprotinin was purchased from Fisher Bioreagents (New Zealand). Sodium dodecyl sulfate (SDS), Tris-HCl, Tris-base, glycine, methanol, Tween-20, magnesium chloride, ethylenediaminetetraacetic acid disodium salt dihydrate, dithiothreitol, and glacial acetic acid were purchased from Fisher Scientific (Pittsburgh, PA). N,N,N’,N’- tetramethyl ethylenediamine and glycylglycine were purchased from Acros Organics (New Jersey, USA). Protogel 30% acrylamide: 0.8% (w/v) bis-acrylamide stock solution was obtained from National Diagnostics, Atlanta, GA. Adenosine-5’-triphosphate disodium salt hydrate and coenzyme A trilithium salt dihydrate were procured from MP Biomedicals, LLC (Illkirch, France). MitoTracker Deep Red FM and DMSO were purchased from Life Technologies, (Carlsbad, CA). Calcein-AM was purchased from BD Pharmingen BD Biosciences (San Jose, CA). Branched polyethyleneimine (PEI, molecular mass ∼ 25 kD) was purchased from Sigma-Aldrich, Saint Louis, MO. Chemicals used in the Seahorse experiments were purchased from Sigma-Aldrich (St. Louis, MO). All other chemicals used were of analytical or cell culture grade and were used as received from the manufacturers.

#### Kits

Pierce Bicinchoninic acid (BCA) and MicroBCA protein assay kits were purchased from Thermo Scientific (Rockford, IL). Quant-iT PicoGreen dsDNA assay kit was procured from Molecular Probes, Inc. (Eugene, OR). CellTiter-Glo-luminescent viability assay reagent was procured from Promega (Madison, WI).

#### Antibodies

Primary mouse antibody to ATP5A (MW 53 kD) (Catalog #ab14748) and rabbit GAPDH polyclonal antibody (MW 36 kD) (Catalog #ab8245) were purchased from Abcam (Cambridge, MA). Alexa Fluor 790-conjugated AffiniPure Donkey Anti Mouse IgG (H+L) was purchased from Jackson ImmunoResearch Lab Inc. (West Grove, PA).

### 2.2. Cell lines and culture

Immortalized human cerebral microvascular endothelial cell line (hCMEC/D3) was purchased from Cedarlane Laboratories, Burlington, Ontario, Canada (Lot #*1x* 102114.3C-P25). Mouse macrophage cells (ATCC TIB-71, RAW 264.7, Mus musculus) were purchased from ATCC (Manassas, VA).

hCMEC/D3 cells were cultured on well plates or tissue culture flasks pre-coated with 150μg/mL of collagen (Type 1 Rat collagen fibrillar collagen, 90%, Corning, Discovery Labware Inc., Bedford, MA) diluted in 0.02 N acetic acid. The collagen solution was applied to the tissue culture flasks or well plates and the culture ware was incubated for 1 h at 37 °C in a humidified incubator. After 1 h, the collagen solution was removed and the surfaces were rinsed with sterile PBS (1*x*, 0.067 M, HyClone, Logan, UT). The hCMEC/D3 cells were cultured in complete media composed of endothelial cell basal medium-2 (Lonza Walkersville Inc., MD), supplemented with 5% heat-inactivated fetal bovine serum (FBS, Hyclone Laboratories, Logan, UT), 10 mM HEPES (pH 7.4) (Fisher Scientific, Pittsburgh, PA), 100 U/mL-100 μ penicillin-streptomycin (Gibco, Carlsbad , CA), 1% chemically defined lipid concentrate (Sigma-Aldrich, Saint Louis, MO), 5 μg/mL ascorbic acid (Sigma-Aldrich), 1.4 μM hydrocortisone (1μmg/mL, Sigma-Aldrich), and 1 ng/mL of recombinant basic fibroblast growth factor (Sigma-Aldrich). RAW 264.7 macrophages were cultured in complete media composed of high-glucose Dulbecco’s Modified Eagle’s Medium (DMEM (1*x*) containing Glutamax (Gibco, Carlsbad, CA) supplemented with 10% FBS and 1% v/v of penicillin-streptomycin. The culture media was changed after every 2 - 3 days. Conditioned media used for EV isolation was of the same composition except for the lack of FBS to exclude serum-derived EVs.

hCMEC/D3 cells and RAW 264.7 cells were passaged when the cells reached 95-100 % confluency. The hCMEC/D3 cells were detached with Trypsin-EDTA (TrypLE Express 1*x*, Gibco, Denmark) and passaged. The hCMEC/D3 cells were used only between passage numbers 25 and 35. RAW 264.7 cells were passaged by dislodging the adherent cells using a cell scraper.

All cells were maintained at 37 ± 0.5 °C in a humidified 5% CO_2_ incubator (Isotemp, Thermo Fisher Scientific).

### 2.3. Isolation of extracellular vesicles (EVs) from hCMEC/D3 endothelial cells and RAW macrophages

For the generation of EVs, hCMEC/D3 cells were seeded in collagen-coated 175 cm^2^ tissue culture flasks (Greiner Bio-One GmbH, Frickenhausen, Germany) while RAW 264.7 cells were seeded directly in 175 cm^2^ tissue culture flasks containing fresh media and cultured till 90-95% confluency. Upon confluence, the media was carefully removed, and the cell monolayer was gently washed once with 25 mL of pre-warmed sterile PBS and replaced with 25 mL of pre-warmed serum-free media in each flask and cultured in a humidified incubator. After 48 h, the conditioned media was harvested and EVs were isolated by differential centrifugation as described previously [23, 26]. Briefly, the culture supernatant was centrifuged at 300*x*g for 10 min to pellet out the cellular debris. The supernatant was then transferred to fresh tubes and further centrifuged at 2000*x*g for 20 min to pellet out the apoptotic bodies. Following this, the supernatant was carefully collected in polycarbonate tubes (26.3 mL, Beckman Coulter, Indianapolis, IN), and spun at 20,000*x*g for 45 min in a type 50.2Ti rotor (Beckman-Coulter Optima XE-90 ultracentrifuge, Indianapolis, IN) to pellet the microvesicles (MVs). The resulting supernatant from this step was then filtered through a 0.22 μm sterile filter into polycarbonate tubes to remove the larger vesicles and centrifuged at 120,000*x*g for 70 min in a Type 50.2Ti rotor to obtain a pellet of exosomes (EXOs). The pellets containing MVs and EXOs were washed once with sterile PBS and centrifuged again following the procedure described above, i.e., at 20,000*x*g for 45 min and 120,000*x*g for 70 min to isolate the MVs and EXOs, respectively. The entire isolation was conducted at 4 °C. All the EV samples were stored at -80 °C till further use. D3-MVs and –EXOs and RAW-MVs and –EXOs indicate EV subsets obtained from the conditioned media from hCMEC/D3 monolayers and RAW 264.7 cells, respectively. It should be noted our EV isolation allowed isolating EXOs and MVs as separate fractions, but <u>we</u> <u>collectively refer to the cell-secreted, membranous EXOs and MVs as EVs</u> [31]<u>, wherever</u> <u>applicable</u>.

### 2.4. Measurement of EV protein content

The total EV protein content was measured using a MicroBCA protein assay. EV samples were diluted in 1*x* RIPA lysis buffer containing aprotinin (10 μg/mL) and were kept on ice for 15 min to lyse the EVs. A 150 μL volume of each sample was added to an equal volume of the MicroBCA working reagent in a 96 well-plate and incubated for 2 h at 37 °C per the manufacturer’s instructions. Protein concentration was quantified by measuring the absorbance at 562 nm on a microplate reader (Synergy HTX multimode reader, Bio-Tek Instruments Inc.).

### 2.5. Physicochemical characterization of EVs

#### 2.5.1. Particle diameter and Zeta Potential

The particle diameters and zeta potentials of EVs were measured using Zetasizer Nano (Malvern Panalytical Inc., Westborough, PA) at a EV protein concentration of 0.2 - 0.5 mg/mL at a temperature of 25 °C and a scattering angle of 173°. Particle size distribution was measured in 1*x* PBS while zeta potential was measured in 10 mM HEPES buffer, pH 7.4. All measurements were performed in triplicates. The data are represented as mean ± standard deviation (SD) of triplicate measurements.

#### 2.5.2. Protein composition

D3-MVs, D3-EXOs, and hCMEC/D3 cells were lysed using 1*x* RIPA containing aprotinin g/mL) on ice for 30 min. The total protein content was quantified using Micro BCA protein assay. Samples containing 40 μg protein were mixed 3:1 with reducing Laemmli SDS sample buffer, 4*X* (Alfa Aesar, MA), and heated at 95 °C for 5 min on a heating block (Thermo Scientific). The samples and premixed molecular weight protein standards (Precision Plus – 250 kD to 10 kD, Bio-Rad, USA) were electrophoretically separated on a 4-10% gradient gel in Tris-Glycine-SDS buffer at 120V for 90 min. The separated proteins were transferred onto a nitrocellulose membrane (Amersham Protran, 0.45 µm, Germany) at 75 V/300 mA for 90 min. The membrane was washed with Tris-buffered saline-tween 20 buffer (TBS-T buffer, 20 mM Tris-HCl, 150 mM NaCl, and 0.1% Tween-20, pH 7.4) and blocked using Li-Cor Odyssey Blocking buffer at room temperature for 1 h. The membrane was then incubated with ATP5A mouse antibody (1:1000) or and rabbit GAPDH polyclonal antibody (1:2000) prepared in Odyssey blocking solution at 4 °C overnight. The membranes were washed using TBS-T buffer followed by incubation with Alexa Fluor 790 conjugated AffiniPure Donkey Anti Mouse IgG (H+L) (1:30,000) prepared in Odyssey blocking solution at room temperature for 1 h. The membranes were again washed in TBS-T buffer, scanned on an Odyssey infrared imaging system (Li-Cor, Lincoln, NE) equipped with Image Studio 5.2 software.

#### 2.5.3. Membrane integrity of the isolated EVs

We first ran fluorescent sub-micron size reference beads (Invitrogen, Carlsbad, CA) with mean diameters of 20-, 100-, 200-, and 500-nm on an Attune NxT Acoustic Focusing Cytometer (Invitrogen, Singapore). The calibration beads allowed the detection of the MVs and EXOs using a 488/10 nm small particle side scatter filter (Invitrogen, Carlsbad, CA) on the BL1 channel and generate a size reference scale. The voltages for forward (FSC-H) and side (SSC-H) scatters were adjusted to 620 V and 240 V respectively and BL1 intensity was adjusted to 380. A total of 5,000 events were acquired for particles within the 100 – 500 nm gate. Threshold on both forward and side scatter channels were set to 0.1 V. EV samples were run at a flow rate of 25 μL/min. Individual EV samples (approximately 50 μL in a volume containing 40 μg – 60 μg of total EV protein) were labelled with 50 μL of 5 μM Calcein-AM (prepared in 10 mM HEPES, 2.5 mM CaCl_2_, diluted from a reconstituted stock solution of 5 mM Calcein-AM in DMSO) and incubated at 37 °C for 20 min, protected from light and diluted to a final volume of 400 μL in 1*x* PBS immediately before analysis. Control samples included filtered PBS and PBS containing 5 μM calcein AM to rule out any potential noise from the reagent. Triton-X 100 (1 % v/v in PBS) was used to lyse EVs. Freshly isolated EV samples, as well as EV samples stored at different conditions (*a*) frozen at -20 °C for a week, (*b*) samples exposed to three freeze-thaw cycles at -20 °C overnight followed by an ice-thaw for 3 h and (*c*) stored at 4 °C for 3 days, were labelled using calcein to determine EV membrane integrity. The fluorescent signals from the beads and labelled EVs were analyzed using density plots on Attune NxT software.

### 2.6. Pre-transfection of donor cells with Lipofectamine/DNA complexes and isolation of pDNA-loaded EVs

hCMEC/D3 endothelial cells and RAW 264.7 macrophage cells were first transiently transfected with Lipofectamine-pDNA complexes. The donor hCMEC/D3 or RAW 264.7 cells were seeded at 1.5 × 10^6^ cells/well in 24-well plates until 80-100% confluency was achieved.

Lipofectamine 3000-pDNA (Luc-pDNA or GFP-pDNA) complexes or control Lipofectamine alone (no pDNA) were prepared with 1 dose of lipid per manufacturer’s protocol (catalog# L300015, Invitrogen, Carlsbad, CA). The manufacturer’s protocol for Lipofectamine 3000 states that 1.0 µL of P3000 reagent and 0.75 µL of Lipofectamine 3000 reagent correspond to one dose of lipid for a 0.5 µg DNA dose/per well in a 24-well plate setup [32]. Cells were transfected such that the pDNA dose was either 0 μg/well (for the generation of naïve EVs), 0.5 μg/well, or 1.0 μg/well (referred to as Luc0.5 or Luc1.0 henceforth) and incubated for 12 h (the total amount of pDNA transfected per 24-well plate were 0, 12, or 24 μg for the indicated groups). The transfected cells were then cultured in serum-free medium for 48 h post-transfection. The conditioned medium was then pooled from each transfection plate and EVs were isolated from each group as described in section 2.3. D3-Luc-MV and D3-Luc-EXO and RAW-Luc-MV and RAW-Luc-EXO indicate the MVs and EXOs isolated from Luc pDNA transfected hCMEC/D3 and RAW 264.7 cells, respectively. Likewise, D3-MV-GFP and D3-GFP-EXO and RAW-GFP-MV and RAW-GFP-EXO indicate the MVs and EXOs isolated from GFP pDNA transfected hCMEC/D3 and RAW 264.7 cells, respectively. For example, RAW-Luc0-MV, RAW-Luc0.5-MV, and RAW-Luc1.0-MV indicate MVs isolated after transfection of parent cells with 0 μg, 0.5 μg, or 1 μg pDNA per well.

### 2.7. EV protein content in DNA-EVs and quantification of the pDNA content in DNA-EVs

The protein content of the EVs in the DNA-EVs was determined using MicroBCA assay as described in 2.4. The amount of double-stranded DNA in the DNA-EVs was measured using the Quant-iT PicoGreen dsDNA assay kit following the manufacturer’s protocols. DNA-EVs were lysed in 1*x* RIPA buffer containing aprotinin (10 μg/mL) on ice for 30 min followed by dilution in 1*x* TE buffer to a total volume of 100 μL. The volumes of EV suspension were adjusted so that it stayed within the standard curve range (0 μg – 1000 μg of calf thymus DNA/200 μL of assay volume). A 100 μL volume of the diluted Picogreen reagent was added to 100 μL of the EV samples in a black 96-well plate. The contents were mixed in a shaker incubator for 5 min at room temperature in the dark and fluorescence was measured using a microplate reader at excitation and emission wavelengths of 485 and 528 nm, respectively (Synergy HTX multimode reader, Bio-Tek Instruments Inc., USA). The percent of DNA loading in the isolated EVs was calculated using **Equation 1**.

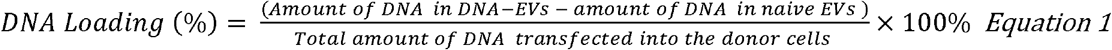

### 2.8. DNA-EV transfection in the recipient brain endothelial cells (BECs)

The transfection activity of Luc DNA-EVs was measured using a luciferase assay. hCMEC/D3 cells were first seeded into collagen-coated 48-well plates at a density of 55,000 cells/well. When the cells reached approximately 80% confluency, the media was freshly replaced with 150 μL of complete medium. D3-Luc-MV; D3-Luc-EXO; RAW-Luc-MV; and RAW-Luc-EXO, each containing 10 ng of DNA were added to each well and incubated for 24 h, 48 h, and 72 h. Untreated hCMEC/D3 cells and cells transfected with Lipofectamine complexes containing 10 ng of Luc-pDNA were used as controls. At 24, 48, and 72 h post-transfection with Luc-EVs, the transfection medium was discarded. The cells were washed with PBS followed by the addition of 100 μL of 1*x* Luciferase cell culture lysis reagent in each well and mixed thoroughly on a shaker for 20 min at room temperature followed by four freeze-thaw cycles (-80 °C for 60 min and 4 °C for 60 min) to completely lyse the cells. The luciferase assay was conducted as previously described by us [26, 33]. The total cellular protein content was quantified using a BCA protein assay using the manufacturer’s protocols. Luciferase expression was expressed as relative light units normalized to total cellular protein (RLU/mg protein). The RLU/mg of protein content for each group was further normalized to untreated cells.

### 2.9. Proteomics analysis of EVs

EVs (25 μg) were electrophoretically separated on 4%-10% Tris-HCl gel. The proteins were visualized using a Bio-Safe Coomassie Blue G-250 stain. The gel lanes were cut into 0.5 mm × 5 mm bands.

#### In-gel trypsin digestion

In-gel trypsin digestion was carried out as previously described [34]. Briefly, gel bands were diced into small pieces (<1 mm^3^) and washed with a solution of 50% acetonitrile/ 25 mM ammonium bicarbonate until visible stain was not present. The gel pieces were then dehydrated with 100% acetonitrile (ACN), reduced with 10 mM dithiothreitol (DTT) at 56 °C for 1 h, then alkylated with 55 mM Iodoacetamide (IAA) at room temperature for 45 min in the dark. Excess DTT and IAA were removed by washing the gel pieces with 25 mM ammonium bicarbonate and then twice with 100% ACN. A solution containing 20 ng/µL sequencing grade modified trypsin (Promega Corporation, Madison, WI; catalog#V511A) and 25 mM ammonium bicarbonate was added to cover the gel pieces and digestion was carried out overnight at 37 °C. The resultant tryptic peptides were extracted from the gel with 70% ACN/5% formic acid (FA), vacuum dried, and reconstituted in 18 µL 0.1% FA for nanoflow liquid-chromatography tandem mass spectrometry (nLC-MS/MS) analysis.

#### Tandem Mass Spectrometry

Tryptic peptides were analyzed by nLC-MS/MS using a NanoAcquity UPLC (Waters’ Corporation, Milford, MA) interfaced to a Velos Pro linear ion trap mass spectrometer (Thermo Fisher Scientific, Waltham, MA). For each analysis, a 1 µL volume of protein digest was injected onto a C18 column (PicoChip column packed with Reprosil C18 3μ chromatography media in a 10.5 cm long, 75 μm ID column with a 15 μm tip, tip, New Objective, Inc., Woburn, MA) and then eluted off to the mass spectrometer using a 37-minute linear gradient of 3-35% ACN/0.1% FA at a flow rate of 300 nL/min.

The Velos Pro was operated in positive ionization mode with a spray voltage of 1.95 kV and capillary temperature of 275 °C. The acquisition consisted of cycles of one full-scan MS1 (AGC of 3x10^4^, 75 ms maximum ion accumulation time, and m/z range of 375-1800) followed by eight MS/MS spectra recorded sequentially for the most abundant ions in the ion trap (minimum signal required 1000 counts, 1x10^4^ AGC target, 100 ms maximum injection time, isolation width 2 m/z, normalized collision energy 35, and activation time 10 ms). Dynamic exclusion (30 s) was enabled to minimize the redundant selection of peptides previously selected for MS/MS.

#### Data Analysis

Collected MS/MS spectra were searched using the MASCOT search engine v2.4.0 (Matrix Science Ltd., London, England) [35] against a Swissprot Homo sapiens database (downloaded on 04/25/2019; 42439 entries) for the human samples and a Uniprot *Mus musculus* database (downloaded 01/19/2019; 94376 entries) for mouse samples. The mass tolerance was set to 1.4 Da for the precursor ions and 0.8 Da for the fragment ions. Peptide identifications were filtered using the PeptideProphet and ProteinProphet algorithms with a protein threshold cut-off of 99%, minimum of 2 peptides, and peptide threshold cut-off of 90% implemented in Scaffold v4.11.0 (Proteome software, Portland, OR).

### 2.10. Gene ontology and pathway enrichment analysis of EVs

The web service Enrichr [36] was used to conduct enrichment analysis of the D3-EV and RAW-EV gene sets derived from Vesiclepedia (**Table S1**). Gene Ontology (GO), a system to associate collections of genes with biological terms and functions [37, 38], was derived for both the D3-EVs and RAW-EVs through the GO Biological Process database. GO terms were ranked by p-value, calculated using Fisher’s exact test, based on the input gene set. In order to gain mechanistic insights pertaining to the two gene sets of interest, pathway analysis was also conducted using the BioPlanet 2019 database [39] for RAW-EVs and the KEGG 2021 Human database [40–42] for D3-EVs, with outputs being similarly ranked. It should be noted that human-specific databases were used for both the human D3-EVs and mice RAW-EVs as mice-specific databases are not available on Enrichr.

**Table 1.**
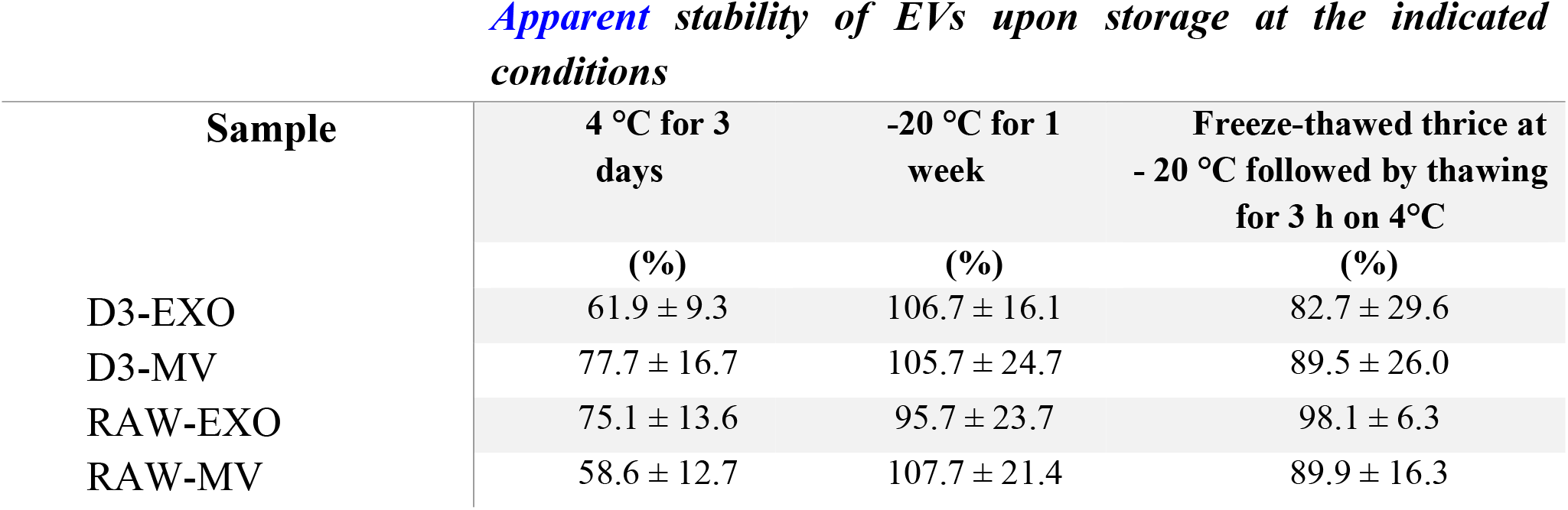
Apparent stability of EVs upon storage at different conditions

| Sample | 4 °C for 3 days | -20 °C for 1 week | Freeze-thawed thrice at - 20 °C followed by thawing for 3 h on 4°C |
| --- | --- | --- | --- |
|  | (%) | (%) | (%) |
| D3-EXO | 61.9 ± 9.3 | 106.7 ± 16.1 | 82.7 ± 29.6 |
| D3-MV | 77.7 ± 16.7 | 105.7 ± 24.7 | 89.5 ± 26.0 |
| RAW-EXO | 75.1 ± 13.6 | 95.7 ± 23.7 | 98.1 ± 6.3 |
| RAW-MV | 58.6 ± 12.7 | 107.7 ± 21.4 | 89.9 ± 16.3 |

### 2.11. Mitochondrial transfer from EVs to the recipient brain endothelial cells

#### 2.11.1. Isolation of EVs from source/donor cells pre-labelled with MitoTracker Deep Red FM

Confluent 175 cm^2^ tissue culture flasks containing hCMEC/D3 cells or RAW 264.7 cells were washed with pre-warmed PBS followed by incubation with 250 nM of MitoTracker Deep Red FM (MitoT) (diluted in the respective growth medium) for 30 min at 37 °C in the dark. The dye-containing medium was then replaced with their respective complete medium and further incubated at 37 °C for 1 h, followed by washing with PBS and incubation with serum-free medium for 16 h. The conditioned media was then subjected to ultracentrifugation as described in section 2.3 to isolate EVs. The isolated pellets were resuspended in 1 mL of sterile PBS. D3- MitoT-MV and D3-MitoT-EXO and RAW-MitoT-MV and RAW-MitoT-EXO indicate the MVs and EXOs isolated from MitoT-labelled hCMEC/D3 and RAW 264.7 cells, respectively. The protein content in MitoT-EVs was determined using a MicroBCA assay.

#### 2.11.2. Treatment of the recipient endothelial cells using MitoT-EVs

Confluent 48-well plates of hCMEC/D3 cells were prepared as described earlier. The cells were then incubated with D3-MitoT-MV, D3-MitoT-EXO, RAW-MitoT-MV, and RAW-MitoT-EXO at different EV protein amounts viz., 3, 24, 100, and 600 μg per well. The plates were incubated for 24, 48, and 72 h at 37 °C in the dark. After incubation, the media was replaced with a phenol red-free DMEM medium. The cells were observed under an Olympus IX 73 epifluorescent inverted microscope (Olympus, Pittsburgh, PA) to detect MitoT signals using the Cyanine-5 (C) channel (Cy5, excitation 635/18 nm and emission 692/40 nm) and under phase contrast settings at 20x magnification. The images were processed using cellSens dimension software (Olympus, USA) and Image J software (NIH). Cell monolayers stained with 250 nM MitoT for 30 minutes in the dark were used as a positive control to detect MitoT signals and unstained cells were used as an additional control. Image contrast was adjusted using ImageJ (NIH). Image analysis was performed by measuring mean intensity using *ImageJ* software. Each image underwent auto-thresholding with identical parameters. Mean intensity values in the treated slices were normalized to control, untreated slices. Statistical analysis was performed using repeated measures one-way ANOVA using GraphPad Prism 9.1.2 software.

#### 2.11.3. Mitochondrial Functional Assessment

Oxygen consumption rate (OCR) and extracellular acidification rate (ECAR) were measured in hypoxic hCMEC/D3 monolayers (20,000 cells/well cultured for four days) in a Seahorse XF96 plate by XF analysis (XF24, Agilent Seahorse Technologies) as previously described [43]. Cells were treated using EXOs or MVs at doses of 30 or 150 µg EV protein/cm^2^ well area corresponding to doses of 3.4 or 17.1 µg EV protein/well in the Seahorse XF96 plate. After measurement of basal OCR, 2.5 μmol/L oligomycin A (proton leak), 0.7 μmol/L FCCP (to measure maximal OCR) were consecutively added. Basal glycolytic rate was calculated by determining the ECAR that is sensitive to 2-DG (100 mmol/L). The assay was performed in non-buffered Dulbecco’s modified Eagle medium supplemented with 25 mmol/L glucose, 1 mM pyruvate, and 2 mmol/L glutamine. All rates were normalized to cellular protein content measured using MicroBCA assay. Data reported indicate average ± SEM from three wells.

#### 2.11.4. Uptake of MitoT-EVs into mice acute brain slices

##### Acute Brain Slice Preparation

The Institutional Animal Care and Use Committee (IACUC) at the University of Colorado approved all experimental protocols in accordance with the National Institutes of Health and guidelines for the care and use of animals in research. Adult (20–25 g) male C57Bl/6 (8–12 weeks) mice purchased from Charles River Laboratory (Fredrickson, NC) were used for this study. All mice were housed in standard 12-h light dark cycle with free access to food and water.

All experiments in the study adhered to the ARRIVE guidelines for animal experiments. Following middle cerebral artery occlusion sham surgery [44–46], mice were anesthetized with isoflurane (3%) and transcardially perfused with artificial cerebral spinal fluid (ACSF: 126 mmol/L NaCl, 2.5 mmol/L KCl, 25 mmol/L NaHCO_3_, 1.3 mmol/L NaH_2_PO_4_, 2.5 mmol/L CaCl_2_, 1.2 mmol/L MgCl_2,_ and 12 mmol/L glucose, pH 7.4) oxygenated with 95% O_2_/5% CO_2_ and at ice-cold conditions (2–5) for 2 min before decapitation. Brains were removed and horizontal cortical or hippocampal sections (300 μM thick) were cut in ice-cold ACSF using a VT1200S Vibratome (Leica, Buffalo Grove, IL, USA) and were recovered in ACSF for 30 minutes at 37 °C before treatment with MitoT-EVs.

##### Incubation with MitoT-EVs

Acute brain slices were incubated in 50 µg/mL of D3-MitoT-EXO and D3-MitoT-MV diluted in normal ACSF for 2 h at 37°C, and counterstained using Hoechst 33258. Non-incubated slices were used as a negative control. The slices were fixed in 4 % of paraformaldehyde overnight at 4°C and were washed in PBS prior to mounting. The slices were imaged using a confocal microscope (Olympus FV1000 laser scanning confocal microscope) equipped with an Olympus Fluoview imaging software (Center Valley, PA, USA) under the cyanine-5 channel (Cy5, excitation 651 nm and emission 670 nm) for visualizing MitoT signals. Image analysis was performed by measuring mean intensity using *ImageJ* software. Each image underwent auto-thresholding with identical parameters. Statistical differences were analysed using repeated measures one-way ANOVA using GraphPad Prism 9.1.2 software.

### 2.12. Effects of EV exposure on the relative cellular ATP levels of the recipient BECs

#### 2.12.1. Effect on normoxic BEC cultures

We determined the effects of EV exposure using the CellTiter-Glo luminescent cell viability assay following the manufacturer’s protocol. Briefly, hCMEC/D3 cells (16,000/well) were seeded in 96-well plates for 24 h in 200 μL of D3 complete media at 37 °C and 5% CO_2_ in a humidified incubator. After confluency, the complete media was replaced with fresh media containing EVs at different protein doses (in a total volume 100 μL/well) and incubated for 72 h. The resulting ATP/cell viability levels in each group was measured as discussed in the following section 2.12.2.

#### 2.12.2. Effect on hypoxic BEC cultures

BECs were exposed to oxygen-glucose deprivation (OGD) as follows: confluent hCMEC/D3 monolayers were washed with pre-warmed PBS, and replaced by glucose-free media as described in [47] and placed in a hypoxia chamber (Billups-Rothenberg, CA, USA) saturated with a 5-7 min flush of 90% N_2_, 5% H_2_, 5% CO_2_ (25 l/min). The sealed hypoxic chamber was kept at 37 °C in a humidified incubator. Different periods of OGD exposure were evaluated to induce endothelial cell death. After exposure of the 96-well plates to the optimized OGD time of 4 h, the media was replaced with 100 μL of OGD medium containing EVs suspended in PBS containing varying amounts of total EV protein and incubated in normoxic conditions (in a humidified 5% CO_2_ incubator) for the indicated times. Healthy cells (non-OGD) cultured under normoxic conditions (∼100% viability) and OGD-exposed cells subsequently cultured in normoxic conditions (∼0% viability) were used as controls. Post-treatment with EVs, cells were washed with pre-warmed PBS followed by the addition of 60 μL of complete growth medium and 60 μL of CellTiter-Glo 2.0 assay reagent to each well. The wells were incubated in a shaker at room temperature for 15 min in the dark. After 15 min, 60 μ of the solution from each well were aliquoted into an opaque, white 96-well plate luminescence plate (Fisherbrand). Relative luminescent signals were measured using Synergy HTX multimode reader (Bio-Tek Instruments Inc., USA) at 1 sec integration time. The relative ATP levels (%) was calculated after normalizing the relative luminescence units (RLU) of treated cells to those untreated cells as shown in **equation 2**.

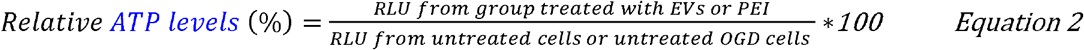

### 2.13. Formation of EV-ATP5A protein complexes

Complexes of recombinant ATP5A1 protein were formed with the EVs by mixing naïve EVs (EXOs/MV 1:1 mixture) and ATP5A at an EV:ATP5A protein weight/weight (w/w) ratio of 5:1. Precalculated volumes of ATP5A solution were added slowly along the walls of the microtube containing EVs diluted in 1*x* PBS. The complexes were prepared by mixing these solutions and vortexed at a setting of ‘5’ on a vortex mixer (Fisher Analog Vortex, 120V, Fisher Scientific, USA) for 30 secs. After mixing, the tubes containing the complexes were spun down for five seconds and allowed to stand at room temperature for 30 min prior to use in experiments.

#### 2.13.1. Native gel electrophoresis

The formation of the EV-ATP5A complexes was confirmed by native polyacrylamide gel electrophoresis. Free ATP5A protein, naïve EV samples, or EV/ATP5A complexes were mixed with an equal volume of native sample buffer (BioRad) and resolved on 4-10% gel in 25 mM Tris, 192 mM Glycine, pH 8.3 at 100 V for 2 h. The gels were then stained with Bio-safe Coomassie Stain solution overnight before scanning using an Odyssey imager (LI-COR Inc., Lincoln, NE) at the 800 nm near-infrared channel.

#### 2.13.2. Effect of the exposure of EV/ATP5A complexes on the ATP levels in the recipient endothelial cells

ATP5A doses of 100, 200, and 300 ng per 0.32 cm^2^ of 96-well plate were used in these studies. The EV-ATP5A complexes were diluted in OGD media before addition to cells 4 h post-OGD exposure. The cells were incubated with the indicated samples for 4 h, washed with pre-warmed PBS, and resulting ATP levels were determined by CellTiter Glo assay (as described in section 2.11.2). The effects of the treatment were expressed as the resulting ATP levels compared to the untreated OGD cells subjected to reoxygenated/normoxic conditions for 4 h. Relative luminescent signals were measured using Synergy HTX multimode reader (Bio-Tek Instruments Inc., USA) at 1 sec integration time. The relative ATP levels (%) was calculated after normalizing the relative luminescence units (RLU) of treated cells to those untreated cells as shown in **equation 3**.

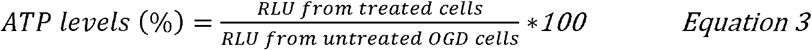

### 2.14. Statistical analysis

The number of independent experiments for each study is indicated in each figure or table legend. Each experiment was repeated at least thrice to confirm data reproducibility. The results are expressed as mean ± standard deviation (SD). Comparative analyses were performed using either one-way or two-way ANOVA using GraphPad Prism v8 (GraphPad Software, San Diego, CA). Wherever applicable, one-way ANOVA was done using Bonferroni’s multiple comparisons test. Alpha was set at 0.05.

## 3. Results and Discussion

The objectives of the current study are two-fold: as described in the introduction section, we first sought out to re-confirm our previous results on the DNA loading and transfection of EVs from two different parent cell sources: brain endothelial cells (BECs) and macrophages, to eliminate any operator-induced systematic biases. In this study, we also conducted proteomic and gene ontology and pathway enrichment analysis to understand possible differences between the BEC- *vs*. RAW-derived EVs. The second goal of the study was to determine the potential effects of the innate EV mitochondrial load on the cellular energetics in the recipient ischemic endothelial cells. In our previous study, we surprisingly observed a higher Luc-DNA loading in the EVs derived from RAW 264.7 macrophages (when the cells were pre-transfected with a lower 0.5 µg DNA/well compared to a higher 1.0 µg DNA/well dose) compared to BEC-derived EVs [26]. In addition, we also observed a higher DNA loading in the smaller exosomes (EXOs) compared to the larger microvesicles (MVs)—this finding was in direct contrast to the findings previously by Kanada *et al.* who reported that while EXOs failed to show expression of the reporter protein encoded by the exogenous pDNA, the larger MVs showed transfer and subsequent expression pDNA-encoded reporter protein in the recipient HEK293FT cells [23]. To rule out any unintentional, operator-induced biases in the experimental regime, an independent operator repeated the donor cell transfection and isolated DNA-loaded EVs from the brain endothelial and macrophage cell lines. We compared the physicochemical characteristics of the DNA-loaded EVs and their transfection activity in the recipient endothelial cells.

The brain endothelial cells have higher metabolic activity compared to the other non-brain endothelial cells [48]. Under ischemic/hypoxic conditions, the endothelial cells are susceptible to undergo apoptosis resulting in increased mitochondrial dependence for metabolism and survival [49]. During the biogenesis of EVs, mitochondria, mitochondrial proteins, or mitochondrial DNA are incorporated into these vesicles and can be transferred between cells [27]. Depolarized mitochondria can be transferred to cells under oxidative stress via MVs resulting in enhanced bioenergetics and cell survival of the recipient cells [27]. Mouse bone marrow-derived stromal cells derived MVs transferred mitochondria and increased the alveolar ATP levels in lipopolysaccharide-exposed mice lungs [29]. Airway myeloid-derived regulatory cells-derived EXOs have also been reported to transfer mitochondria to T cells and localize with the T cell’s mitochondrial network [28].

The second objective of our work is to determine whether the EV mitochondrial load can be transferred to ischemic endothelial cells in an *in vitro* oxygen-glucose deprivation model of stroke and to neurons in mice acute brain cortical and hippocampal slices. The lack of ATP serves as the initial trigger in the ionic, biochemical, and cellular events that cause death and damage during stroke [50]. Therefore, we, hypothesized that the transfer of innate EV mitochondria may switch the endothelial cell fate from death to survival. The underlying rationale for our hypothesis is that protecting the BECs lining the BBB will subsequently allow it to maintain its barrier properties and limit BBB dysfunction-induced neurological damage in diseases like stroke.

### 3.1. Isolation and characterization of hCMEC/D3 endothelial cell- and RAW 264.7 macrophage-derived EVs

As previously reported by us [26], a cell model of the human BBB, hCMEC/D3 endothelial cells, and RAW 264.7 macrophages were used to compare if the donor/source cell line has any effects on the extent of DNA loading for subsequent transfection into the recipient endothelial cells. Naïve EVs isolated from hCMEC/D3 endothelial and RAW 264.7 macrophage cells showed average particle diameters ranging between 100 to 250 nm as shown in **Fig. 1b-e**. We noted a significant difference among the average particle diameters for EXOs and MVs isolated from both cell sources. D3-EXO, D3-MV, and RAW-EXO showed heterogeneous size distributions as expected for cell-derived vesicles. A smaller fraction of EXO sized <100 nm was seen in D3-EXO while D3-MV and RAW-EXO showed particle populations >150 nm. MVs are known to be heterogeneous in size with effective particle diameters ranging from 200 - 1500 nm [51, 52]. The overall sizes of EXOs and MVs isolated from both cell lines were in agreement with previous studies [23]. However, EVs are also prone to aggregation, which may have resulted in particles sizes >150 nm. These size ranges suggest the increased likelihood of EVs entering recipient cells via endocytosis [53]. The zeta potentials of EV samples ranged between - 4 and -12 mV (**Fig. 1a**). The negative zeta potential is attributed to their anionic membrane lipids like phosphatidylserine and glycosylated proteins that are incorporated into the EXOs and MVs during their biogenesis [54, 55].

**Figure 1.**
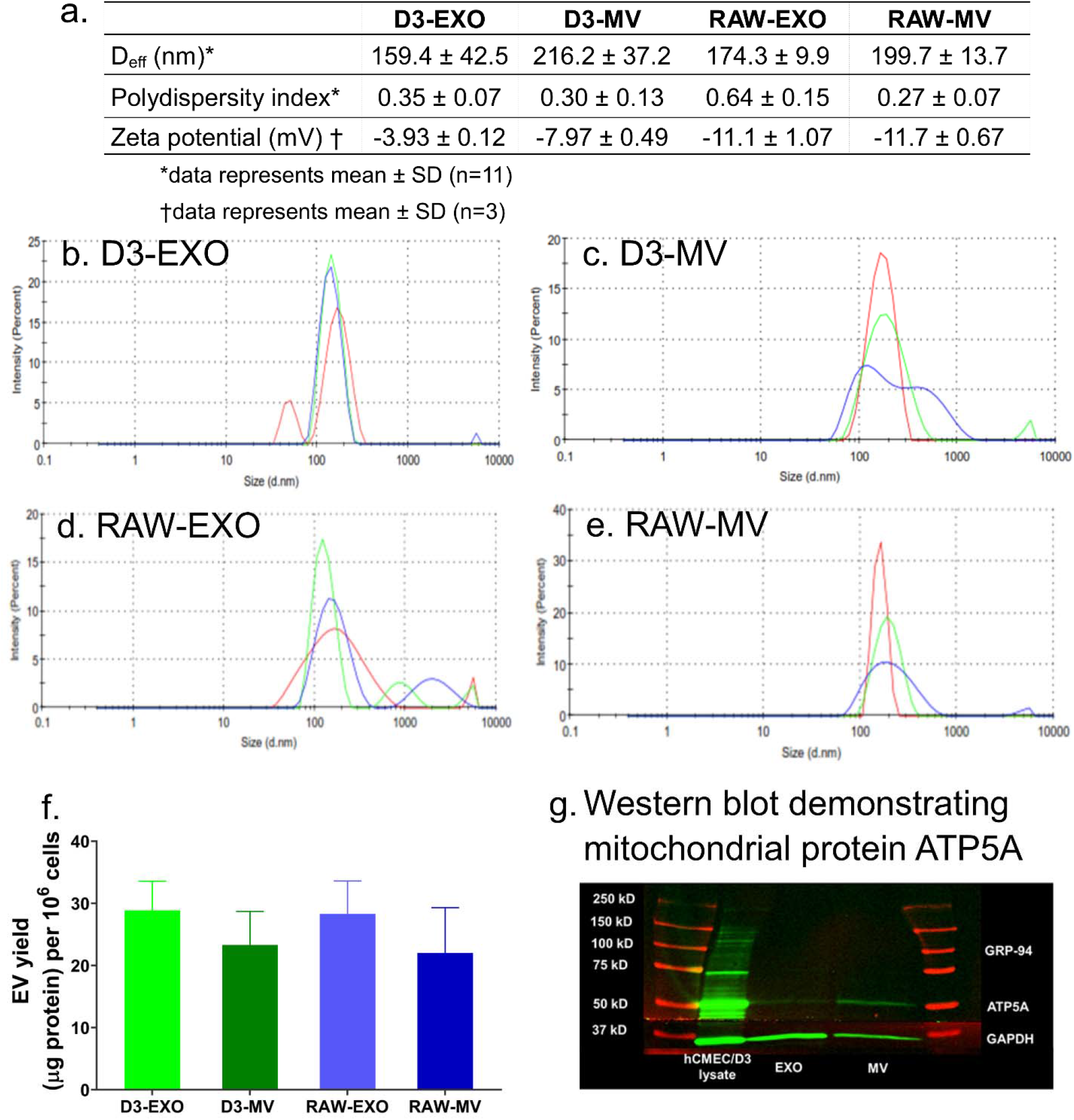
Characterization of EVs derived from hCMEC/D3 (D3-EXO and D3-MV) and RAW 264.7 cells (RAW-EXO and RAW-MV). (*a*) Physicochemical characteristics of EVs: Effective particle diameter (D_eff_), polydispersity index, and zeta potential were measured using dynamic light scattering (DLS). The samples at a protein concentration of 0.2 – 0.5 mg/mL were resuspended in 1*x* PBS and 10 mM HEPES buffer, pH 7.4 for D_eff_ and zeta potential measurements, respectively. Representative DLS intensity plots of (*b*) D3-EXO (*c*) D3-MV (*d*) RAW-EXO and (*e*) RAW-MV obtained from measurements on a Malvern Nano Zetasizer. The different traces indicate three measurements of the same sample. (*f*) EV yield normalized to per million cells of hCMEC/D3 or RAW 264.7 *(g)* Western blotting to confirm mitochondria-specific EV markers. 25 µg total protein was loaded in a 4-10% SDS gel and electrophoresed at 120 V. The separated proteins were transferred on nitrocellulose membrane and stained with ATP5A and GAPDH antibodies. The blots were imaged on Odyssey imager (LI COR Inc., Lincoln, NE) at 800 nm near-infrared channel and processed using ImageStudio 5.2 software.

Normalization of the EV protein yields to the total cell numbers resulted in values of 28.9±4.7 µg/10^6^ cells and 23.4±5.4 µg/10^6^ cells for D3-EXO and D3-MV, respectively, and 28.3±5.3 µg/10^6^ cells and 22.0±7.3 µg/10^6^ cells for RAW-EXO and RAW-MV, respectively (**Fig. 1f**). It should be noted that we isolate EVs from a “conditioned” medium that lacks serum to avoid collecting serum-derived EXOs. Therefore, it is likely that cells may be sensitive to serum withdrawal. Li *et al.* investigated the release and protein composition of EVs derived from cells cultured in medium supplemented with serum-depleted EVs and EVs derived from cells cultured in serum-free media. The authors reported that although serum-free medium induced cellular stress, it also increased the release of EVs along with stress proteins like macrophage migration inhibitory factor and epoxide hydrolase 1 [56]. Contrarily, other reports suggested that the presence of serum growth factors in culture media influences the intracellular calcium that is responsible for vesicle shedding [51, 57, 58].

MVs are known to selectively enriched with the mitochondrial protein ATP5A, a catalytic subunit of ATP synthase [59] compared to the EXO fraction. Our western blotting analysis confirmed the same (**Fig. 1g**). Glyceraldehyde 3-phosphate dehydrogenase (GAPDH) protein was also used as an additional protein in our study. GAPDH (36 kD) is commonly used as an internal control to ensure uniform protein loading. We noticed a lower expression of GAPDH in the MV fraction compared to the EXO fraction. This is consistent with earlier findings that reported a greater GAPDH expression in EXOs than in the MVs [23].

The integrity of EV membranes and its stability upon storage were determined using flow cytometry using previously reported methods [60, 61]. Calcein AM is a non-fluorescent, membrane-permeable dye that undergoes hydrolysis by intravesicular esterases present only in an intact vesicle [62]. Derivatives of calcein especially its acetomethoxy derivative (Calcein-AM) have been extensively studied for live cell tracking and detection. This has also been applied to distinguish functional and metabolically-active EVs from the disrupted EV or protein debris [63]. Fluorescent sub-micron size reference beads with mean diameters of 20-, 100-, 200-, and 500-nm were used to generate a size reference scale. Gating was applied to exclude the background laser noise in the scatter plots [63]. The detection gate was set between 100 and 500 nm to allow the detection of the beads alone (**Fig. S1a,b**). Gated events for EVs (R1) corresponding to the free calcein-AM solution in 1*x* PBS (**Fig. S1c,d**) and non-stained EVs (**Fig. S1e-h**) were used as controls. Additionally, an EV sample lysed using 1% (v/v) of Triton X-100 was used as a negative control. As expected, no positive events for EVs were observed following the treatment of EVs with Triton X-100 (**Fig. S1m-p**).

RAW-MV (**Fig. S1i**), RAW-EXO (**Fig. S1j**), D3-MV (**Fig. S1k**), D3-EXO (**Fig. S1l**) with calcein-positive counts were noted to be present corresponding to particle diameters ranging from 100 to 200 nm. This correlates with our DLS data wherein the mean effective particle size diameters of D3-EXO and D3-MVs ranged from 160 to 250 nm. Lucchetti *et al.* also reported 100 nm-sized EXOs with a refractive index less than 1.4 based on the side scatter signal intensity compared to 100 nm polystyrene beads (refractive index of ∼1.6) [63].

To investigate the effect of storage conditions on the integrity of EV membranes, EV samples were stored at -20 °C and 4 °C for different periods until analysis and compared to the control group of freshly-isolated EV samples. From the SSC/FSC scatter profile, it appeared that their sizes were comparable to the control, but the number of EV events was significantly altered. After storage for 3 days at 4 °C, the number of EV events for D3-EXO was markedly lower compared to the control (**Table 1**). Freezing the EVs at -20 °C retained the number of EV events better than those samples that were subjected to three freeze-thaw cycles (frozen thrice at -20 °C followed by thawing for 3 h at 4°C). The apparent stability of EVs as a function of their membrane integrity was calculated using **equation 4**.

Apparent stability of EVs at the test storage condition (%) was calculated using the following equation:

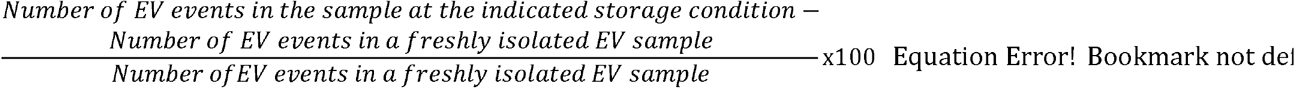

It can be noted that in some samples, the number of EVs events were higher than the control sample. A similar observation was seen earlier in [61], wherein the number of MV events were higher upon storage at -80 °C with a simultaneous decrease in the size of MVs, the mechanism for which is currently unknown. Similarly, a variation and a trend towards an increase in EV count were observed upon storage at – 20 °C and – 80 °C by Jeyaram and colleagues [64]. Likewise, Kong *et al.* observed that MVs isolated from EDTA-anticoagulated plasma stored at - 80 °C for 4 weeks increased the MV count and gradually decreased the MV size. The number of MVs in plasma increased almost two-fold upon storage for 4 weeks at -80 °C versus the control sample which was collected and centrifuged immediately without any storage. The exact mechanism for the apparent “increase” in EV/particle numbers is currently unknown [61]. Lőrincz *et al.* observed an increase in the geometric mean of the flow cytometer side-scatter distribution for EV fractions after storage for 28 days at -20 °C [65]. This could be due to a shift in the EV size upon storage at -20 °C [64] likely due to deaggregation/structural change of the EVs [65]. Lőrincz *et al.* suggested that the swelling of vesicles upon storage may have caused more of the smaller-sized vesicle fraction to be detected in the scatter plots [65]. The apparent stability of EV is calculated by the difference in the number of events recorded in the EV samples stored at different storage conditions in comparison to freshly isolated EV samples. Therefore, an increase in the particle counts in the gated area is subsequently reflected as increased “apparent” stability values.

### 3.2. DNA-loaded EVs derived from hCMEC/D3 endothelial cells and RAW 264.7 macrophages

As previously reported by us [26], a passive transfection-based approach was used to engineer the EVs using Luc- pDNA as a model plasmid. We used Lipofectamine:pDNA ratios of 1:1 to transfect the parent cells to avoid any side effects of excess/free lipofectamine. The non-specific toxicity of cationic lipids [66], their potential to alter the cell gene expression, and their potential effects to alter the innate EV cargo [67] cannot be ignored. Therefore, we used the minimally-required amount of lipofectamine to transfect pDNA into the parent cells while varying the amount of Luc DNA – 0.5 or 1.0 µg (Luc0.5 or Luc1.0), transfected into the parent cells. We used a QuantiT Picogreen assay to determine the dsDNA loading in the EVs isolated from hCMEC/D3 endothelial cells and RAW 264.7 macrophages.

Consistent with our previous findings, our data showed that the RAW-Luc-EXO loaded more DNA than the D3-Luc-EXO (*p < 0.001* for Luc0.5) compared to the RAW-Luc-MV and D3- Luc-MV. Moreover, RAW-Luc-MV revealed no significant difference in their DNA loading at both the DNA doses of 0.5 and 1.0 µg/well compared to D3-Luc-MV at their respective doses. However, Luc-pDNA loading in RAW-Luc0.5-MV was significantly lower (*p < 0.0001*) than RAW- Luc0.5-EXO. The percent DNA loading (**Fig. 2a**) was calculated using **equation 1**. Both D3-Luc0.5-EXO and RAW-Luc0.5-EXO showed almost a three-fold increase in the percent DNA loading compared to D3-Luc1.0-EXO and RAW-Luc1.0-EXO. No significant differences were noted in the extent of DNA loading in MVs derived from hCMEC/D3 endothelial and RAW 264.7 macrophage cells. The maximum level of DNA loading was observed in RAW- Luc0.5-EXO (approximately 6.5%) followed by RAW-Luc1.0-EXO and D3-Luc0.5-EXO.

**Figure 2.**
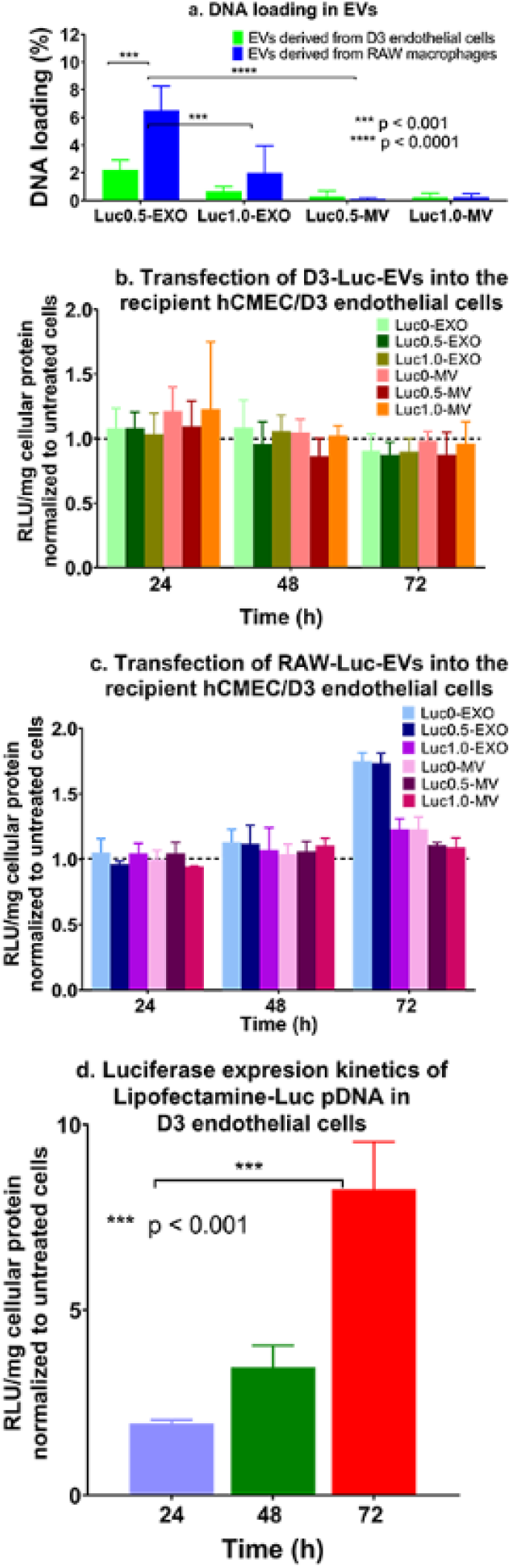
Measurement of DNA content in Luc pDNA-loaded EVs derived from hCMEC/D3 endothelial cells and RAW 264.7 macrophages and their transfection activity in the recipient hCMEC/D3 endothelial cells. hCMEC/D3 endothelial and RAW 264.7 macrophages were transfected with Lipofectamine-Luc pDNA at a pDNA dose of 0.5 μg/ or 1.0 μg/well in a 24–well-plate (n = 3). (*a*) The percent Luc-DNA loading in the isolated EVs was measured by Quant-iT Picogreen dsDNA assay using *Equation 1* (*b*) Transfection of D3-derived Luc EVs, (*c*) RAW-derived Luc-EVs and (*d*) Lipofectamine/Luc pDNA complexes into the recipient hCMEC/D3 endothelial cells at a DNA dose of 10 ng of DNA/well (n = 4). Luciferase gene expression was expressed as Relative light units (RLU) normalized to total cellular protein content and further normalized to values from the control, untreated cells. Data are presented as mean ± SD (n = 4), *\*p < 0.05*, *\*\*p < 0.01*, *\*\*\*p < 0.001*, *\*\*\*\*p < 0.0001* by two-way ANOVA of the indicated groups and Bonferroni’s multiple comparisons test.

The reproducible observations of higher pDNA-Luc loading in RAW-derived EVs may likely be due to the fact that brain endothelial cells possess high intrinsic resistance to transfection compared to the other cell lines even while using the potent non-viral transfection agents [68, 69] and as a result, BEC-derived EVs also concomitantly load lower amounts of the transfected DNA. We also posit that as EXOs are endosomal-derived vesicles compared to the membrane-derived MVs, there is a greater likelihood that the lipofectamine/DNA complexes (that are known to be internalized via endocytosis) or part of the released DNA were entrapped in these EXOs compared to MVs. In other words, there is a natural overlap between the subcellular trafficking of the lipofectamine/DNA complexes and the biogenesis of exosomes. This natural overlap may allow a greater loading of the exogenous DNA into the smaller EXOs compared to the larger MVs that bud off from plasma membranes. Therefore, from two studies conducted by independent operators in our lab ([26] and the current study), we conclude that the smaller EXOs loaded a greater amount of exogenous pDNA compared to the larger MVs. Noteworthy, EV production and release into the culture medium is a dynamic process that depends upon the production rate and the recycling and reuptake rates of EVs back into the cells [70]. This may have resulted in lower pDNA loading in the slow-dividing hCMEC/D3-derived EVs than those derived from rapidly-dividing RAW 264.7 macrophage cells.

Consistent with our previous observations, the loading efficiency of pDNA in the EVs was independent of the pDNA dose transfected into the donor/source cells (0.5 vs. 1 μg/well). An increase in the transfected pDNA dose (Luc1.0-EXO or Luc1.0-MV) did not result in a measurable increase in DNA transfer or the resulting luciferase protein expression compared to Luc0.5-EXO or Luc0.5-MV groups derived from both the cell lines (**Fig. 2b and c**). It has been reported that the loading of DNA into EVs is dependent upon the size of the DNA construct used [25]. Electroporation has shown maximal DNA loading levels of ∼2% for 750 bp DNA. Lamichhane *et al*. reported that linear dsDNA molecules of 250 bp were loaded into the EVs efficiently via electroporation at a maximum amount of 5 μg linear dsDNA [25]. Their data showed that approximately 50 ng of DNA was loaded in 3 × 108 EVs when an initial amount of 2.5 μg of linear dsDNA was used [25]. Plasmid DNA and linear DNA greater than 1000 bp were not loaded using the electroporation approach. The suggested plasmid DNA cut-off is ∼ 4.5 to ∼10 kb albeit the loading efficiencies were < 0.2% [25]. The size of the plasmids used in the study was 6.7 kb and 5.8 kb for Luc-pDNA and GFP-pDNA, respectively. The DNA loading in D3-GFP0.5-EXO and D3-GFP1.0-EXO was 0.58 % and 0.36 % respectively, while that in D3-GFP0.5-MV and D3-GFP1.0-MV were 0.11 % and 0.10 % respectively. As observed in the EVs loaded with Luc-pDNA, the EXOs showed a greater DNA loading capacity compared to MVs. Likewise, the DNA loading in RAW-GFP0.5-EXO and RAW-GFP1.0-EXO was 0.29% and 0.48% respectively, while that in RAW-GFP0.5-MV and RAW-GFP1.0-MV were 0.36% and 0.36% respectively. MVs derived from both the cell lines showed an insignificant gradual increase in exogenous DNA (from 0/naïve EVs to 1 µg/well) and are likely due to the variations in DNA loading that correlates with their biogenesis pathways as discussed in the previous paragraph. Interestingly, our results contradicted previously published reports [23, 25], that suggested that MVs showed a greater potential for transfection compared to EXOs due to increased DNA loading although Lamichhane *et al*. did not observe measurable transfection of the DNA-EVs [25].

The differences in DNA loading between EXOs and MVs may also be explained because EXOs have less endogenous or innate cargo. A proteomics study comparing the protein content of EXOs and MVs reported that MVs are enriched with proteins related to the cytoskeletal network and cortical activity compared to the EXOs [71]. Moreover, if MVs are released through a regulated, generally low, steady-state process as compared to EXOs that are released constitutively [72], the biogenesis of MVs loaded with DNA will also be comparatively lower. No significant changes in the total EV protein content were observed with DNA-loaded EVs and EVs isolated from cells treated with lipofectamine alone (no DNA). This indicated that the biogenesis and release of EVs were not unaffected due to the transfection process using cationic lipids (lipofectamine) at the different pDNA amounts. We further compared the particle diameters of naïve vs. DNA-loaded EVs (**Table 2**) to determine if DNA loading affected EV particle characteristics. We prepared EVs loaded with both luciferase (luc) and eGFP (GFP) pDNA to determine if the type of the pDNA construct affected their particle characteristics. It is well-documented that the physicochemical characteristics of nanoparticle carriers strongly affect their biological activity *in vitro* [33, 73, 74]. We noted that the effective particle diameters of DNA-EVs derived from hCMEC/D3 cells were somewhat unchanged with a slightly higher PdI compared to naïve D3-EVs, however, the DNA-EVs derived from RAW 264.7 cells seemed to have larger vesicular sizes as well as showed a heterogeneous sample with a greater PdI, compared to naïve RAW-EVs. The noted changes in the particle diameters of RAW-EVs may be reflective of the exogenous DNA loading.

**Table 2.** Effective particle diameters (D_eff_) of DNA-EVs

| Sample | Average $D_{\text{eff}}$ (nm) | PdI | Sample | Average $D_{\text{eff}}$ (nm) | PdI |
| --- | --- | --- | --- | --- | --- |
| Naïve D3-EXO | $159.4 \pm 42.5$ | $0.4 \pm 0.1$ | | | |
| D3-Luc0-EXO | $133.7 \pm 19.7$ | $0.4 \pm 0.1$ | D3-GFP0-EXO | $156.6 \pm 7.3$ | $0.4 \pm 0.0$ |
| D3-Luc0.5-EXO | $111.1 \pm 5.7$ | $0.4 \pm 0.1$ | D3-GFP0.5-EXO | $122.2 \pm 5.8$ | $0.5 \pm 0.0$ |
| D3-Luc1.0-EXO | $133.4 \pm 25.4$ | $0.3 \pm 0.1$ | D3-GFP1.0-EXO | $118.8 \pm 0.8$ | $0.4 \pm 0.1$ |
| D3-MV | $216.2 \pm 37.2$ | $0.3 \pm 0.1$ | | | |
| D3-Luc0-MV | $190.2 \pm 13.8$ | $0.3 \pm 0.1$ | D3-GFP0-MV | $339.3 \pm 122.3$ | $0.4 \pm 0.1$ |
| D3-Luc0.5-MV | $306.6 \pm 105.5$ | $0.4 \pm 0.1$ | D3-GFP0.5-MV | $204.3 \pm 17.4$ | $0.4 \pm 0.0$ |
| D3-Luc1.0-MV | $368.5 \pm 37.9$ | $0.5 \pm 0.1$ | D3-GFP1.0-MV | $176.1 \pm 22.3$ | $0.4 \pm 0.0$ |
| RAW-EXO | $174.3 \pm 9.9$ | $0.6 \pm 0.2$ | | | |
| RAW-Luc0-EXO | $483.5 \pm 120.4$ | $0.6 \pm 0.1$ | RAW-GFP0-EXO | $594.7 \pm 204.0$ | $0.5 \pm 0.1$ |
| RAW-Luc0.5-EXO | $163.9 \pm 25.4$ | $0.6 \pm 0.1$ | RAW-GFP0.5-EXO | $385.0 \pm 236.2$ | $0.5 \pm 0.1$ |

| EXO |  |  | EXO |  |  |
| --- | --- | --- | --- | --- | --- |
| RAW-Luc1.0- | 122.0 ± 15.8 | 0.3 ± 0.0 | RAW-GFP1.0- | 556.5 ± 303.8 | 0.6 ± 0.3 |
| EXO |  |  | EXO |  |  |
| Naïve RAW-MV | 199.7 ± 13.7 | 0.3 ± 0.1 |  |  |  |
| RAW-Luc0-MV | 472.1 ± 198.0 | 0.5 ± 0.2 | RAW-GFP0-MV | 276.7 ± 37.7 | 0.5 ± 0.1 |
| RAW-Luc0.5- | 668.0 ± 210.6 | 0.6 ± 0.2 | RAW-GFP0.5- | 288.5 ± 23.8 | 0.4 ± 0.0 |
| MV |  |  | MV |  |  |
| RAW-Luc1.0- | 585.4 ± 37.5 | 0.6 ± 0.0 | RAW-GFP1.0- | 386.3 ± 61.3 | 0.5 ± 0.1 |
| MV |  |  | MV |  |  |

**Table 3:**
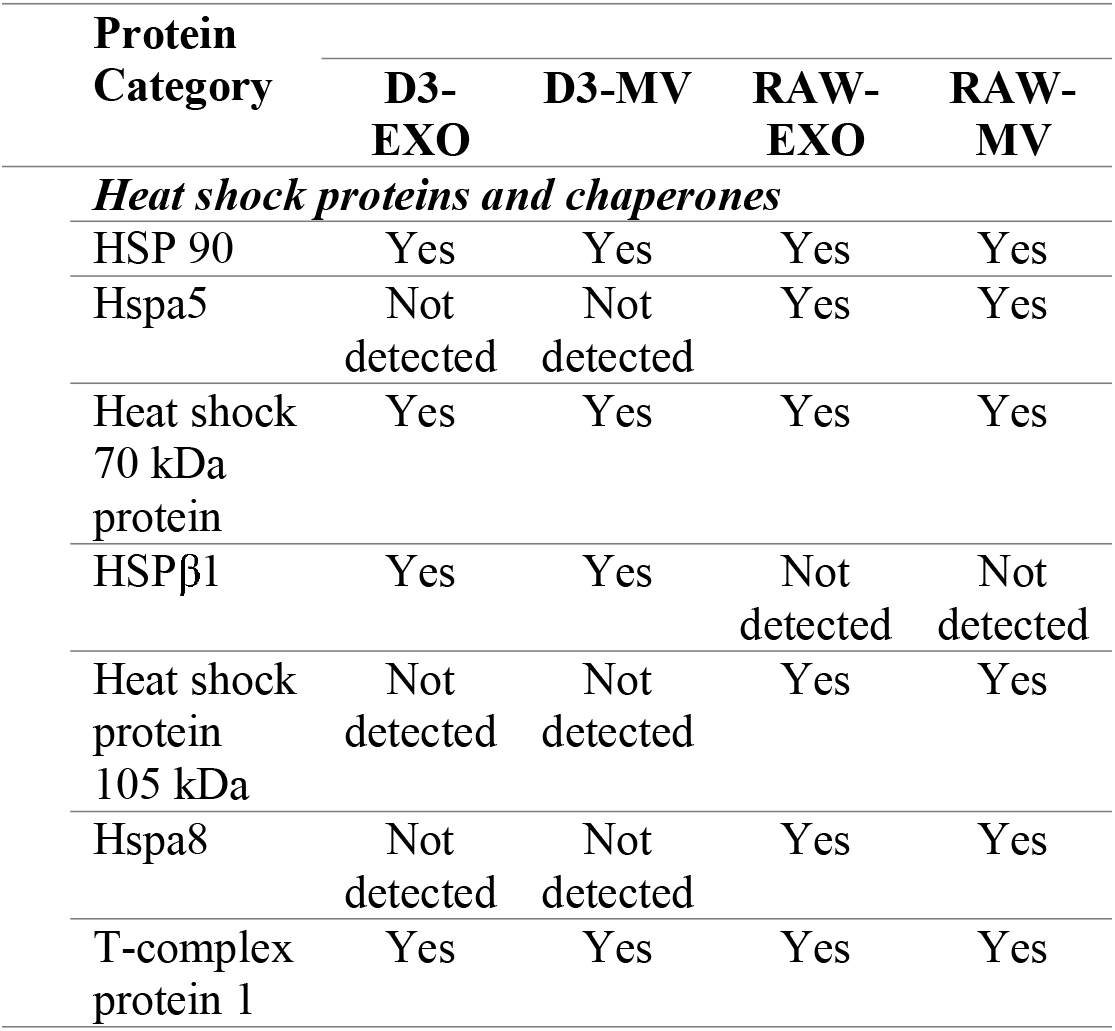
Proteomic analysis highlighting key proteins in EVs (*Yes/not detected indicates the expression of the listed proteins in the respective samples).

### 3.3. Transfection of DNA-loaded EVs in the recipient hCMEC/D3 endothelial cells

We studied the ability of the engineered EVs to transfer Luc-pDNA into recipient D3 endothelial cells *in vitro*. Recipient hCMEC/D3 cells were transfected using Luc-EVs derived from either hCMEC/D3 endothelial cells or RAW macrophages. Transfection activity of D3- derived Luc-EVs is shown in **Fig. 2c** and RAW-derived EVs is shown in **Fig. 2d**. Homotypic EVs with analogous cell membranes typically show a higher cellular uptake due to self-recognition compared to heterotopic EVs [75].

EVs are heterogeneous vesicles and it is known that even EVs obtained from the same cells differ in their content [76]. Willis *et al.* observed that EXOs larger than 80 nm are enriched in flotillin-1 and CD 63, while those smaller than 80 nm are enriched with TSG101 and ALIX [77]. EXO subpopulations are known to exhibit differential packaging of nucleic acids [78]. Given the inherent heterogeneity of these cell-derived vesicles, we decided that the best approach is to transfect the recipient cells at a constant amount of DNA (10 ng), instead of transfecting using a constant number of EVs. Lesson learnt from the non-viral transfection field indicates that amongst factors such as the chemical composition of the carriers (lipids/polymers), carrier/nucleic acid ratios, etc., the DNA dose is one of the critical factors that determine the transfection outcomes.

In comparison to the EVs, the positive control group transfected with Lipofectamine-Luc- pDNA complexes containing an equivalent amount of pDNA showed comparatively higher expression of luciferase (almost 6 to 8-fold increase in luciferase expression compared to untreated cells) and such increases persisted for over 72 h in hCMEC/D3 endothelial cells (**Fig. 2d**). Our findings are consistent with a previous study that reported that cationic lipids showed an increased photon flux on the 2^nd-^day post-transfection that decreased gradually over 7 days [79]. Overall, our results on DNA loading into the EVs were consistent with our previous study. This suggested that any unintentional/operator-induced biases did not cause the higher DNA loading in the smaller EXOs compared to the larger MVs, observations that are in contrast previously published results [23].

### 3.4. Proteomics study to determine potential compositional differences in D3- vs. RAW-derived EVs

We conducted proteomics analyses of the EVs in an effort to understand potential compositional differences between D3- and RAW-derived EVs. Biogenesis of EXOs and MVs involves the packaging of cytoplasmic proteins and membrane proteins in the vesicles. Liquid chromatography-tandem mass spectrometry (LC-MS/MS) was used to qualitatively study the proteins present in EVs. Previous studies have identified 295 proteins in urinary exosomes [80] and 272 in mast cell-derived EXOs [81] upon mapping to Entrez gene identifiers. The proteome profiles of our D3-EVs and RAW-EVs resulted in the identification of 136 proteins in D3-EXOs, 116 proteins in D3-MV, 169 proteins in RAW-EXO, and 141 proteins in RAW-MVs. We compared and studied the overlap of our EV proteins with ‘Vesiclepedia’, a publicly available extracellular vesicle protein database. We identified most of the top 100 proteins enlisted in the Vesiclepedia database (**Table S1**).

**Table S1:** Top 100 proteins identified in the isolated EVs compared to Vesiclepedia database

We speculate that the enrichment of HSP proteins in the EVs may have contributed to the observed ATP increases in the normoxic as well as hypoxic endothelial cultures (**Fig. 6**). Heat-shock 70-kDa proteins (HSP70, HSP71A), constitutive heat-shock proteins (HSPA8 and HSPA5), HSP105, and HSP90 (HSP90A/B) were present in EXOs and MVs isolated from hCMEC/D3 and RAW 264.7 cells. HSP-beta (HSP β1) was specifically expressed in D3-EVs, while HSP105 and HSPA8 (Q3UBA6) were specifically observed in RAW-EVs. HSPA8 and HSP90 are reported to be among the top ten proteins found in most of the EXOs [82].

### 3.5. Gene ontology and pathway enrichment analyses

The gene ID for top-100 proteins identified from the Vesiclepedia data base was further distilled into gene sets exclusively expressed in the D3- and RAW-EVs (**Table S2**).

**Table S2:** Gene IDs for D3-EVs versus RAW-EVs

The GO enrichment analyses for D3-EVs resulted in a very high association to glycolysis-related processes (Fig. 3). Glycolytic process (GO:0006096) was the most overexpressed term, followed by carbohydrate catabolic process (GO:0016052) (p = 6.53 × 10^-11^) and pyruvate metabolic process (GO:0006090) (p = 4.14 × 10^-10^) (Fig. 3a). The GO analyses of RAW-EVs revealed an abundance of neutrophil-related terms, accounting for three out of the four most significant GO terms. Yet, glycolytic process again was the highest ranked (p = 1.60 × 10^-13^), then followed by neutrophil degranulation (GO:0043312) and neutrophil activation involved in immune response (GO:0002283) (p = 5.73 × 10^-13^) (**Fig. 3b**).

**Figure 3.**
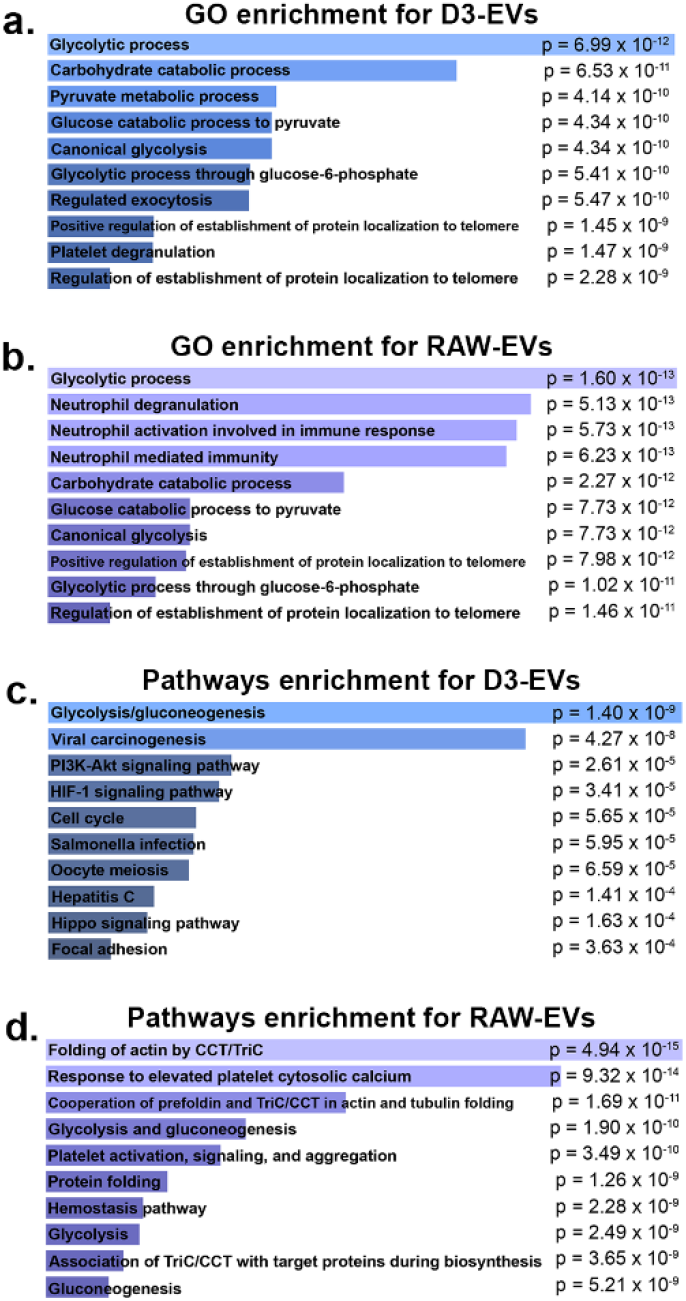
GO and pathway enrichment analyses revealed RAW- and D3-EVs overexpress glycolytic processes. The Enrichr web service was used to analyze enriched pathways and GO terms within the two gene sets for RAW- and D3-EVs. Gene ontology analysis using the 34-gene set for D3- EVs and 42-gene set for RAW-EVs showed that GO terms surrounding glycolytic process are greatly overexpressed in the D3 cells (p = 6.99 × 10^-12^) (**a**), while in addition to glycolytic terms, there were a significant amount of neutrophil-related GO terms (p = 5.13 × 10^-13^) in RAW-EVs (**b**). Pathway analysis results indicated that glycolysis pathways were overexpressed in both D3- and RAW-EVs (p = 1.4 × 10^-9^ and 1.9 × 10^-10^). p-values were calculated using Fisher’s exact test.

Analogous to GO analyses, we derived a list of overexpressed pathways in the two gene sets corresponding to RAW- and D3-EVs to better understand their mechanistic actions. For D3-EVs, glycolysis/gluconeogenesis appeared the most significant (p = 1.40 × 10^-9^), followed by viral carcinogenesis and, after a sizeable gap in significance, PI3K-Akt and HIF-1 signalling pathways (**Fig. 3c**). In RAW-EVs, two of the top three, and three of the top ten, overexpressed pathways were related to the CCT/TriC molecular chaperone (**Fig. 3d**) followed by glycolysis/gluconeogenesis pathways. The results from this enrichment analysis support, for one, the strong metabolic and glycolytic association of D3-EVs. Therefore, we speculated that the D3-derived EVs may be strongly associated with processes that contribute to the mitochondrial transfer (**Fig. 4, 5 and 7**) and those linked with the production of ATP (**Fig. 6**). Interestingly, the analysis also showed numerous overexpressed neutrophil-associated GO terms in RAW-EVs. This can be partially attributed to their immune origin wherein processes like degranulation also contribute to glycolysis, exocytosis, and the production of cell energy [83]. In light of their potent enrichment in glycolytic pathways, we chose to focus on aspects related to EV-mediated mitochondrial transfer for the remainder of this study.

**Figure 4.**
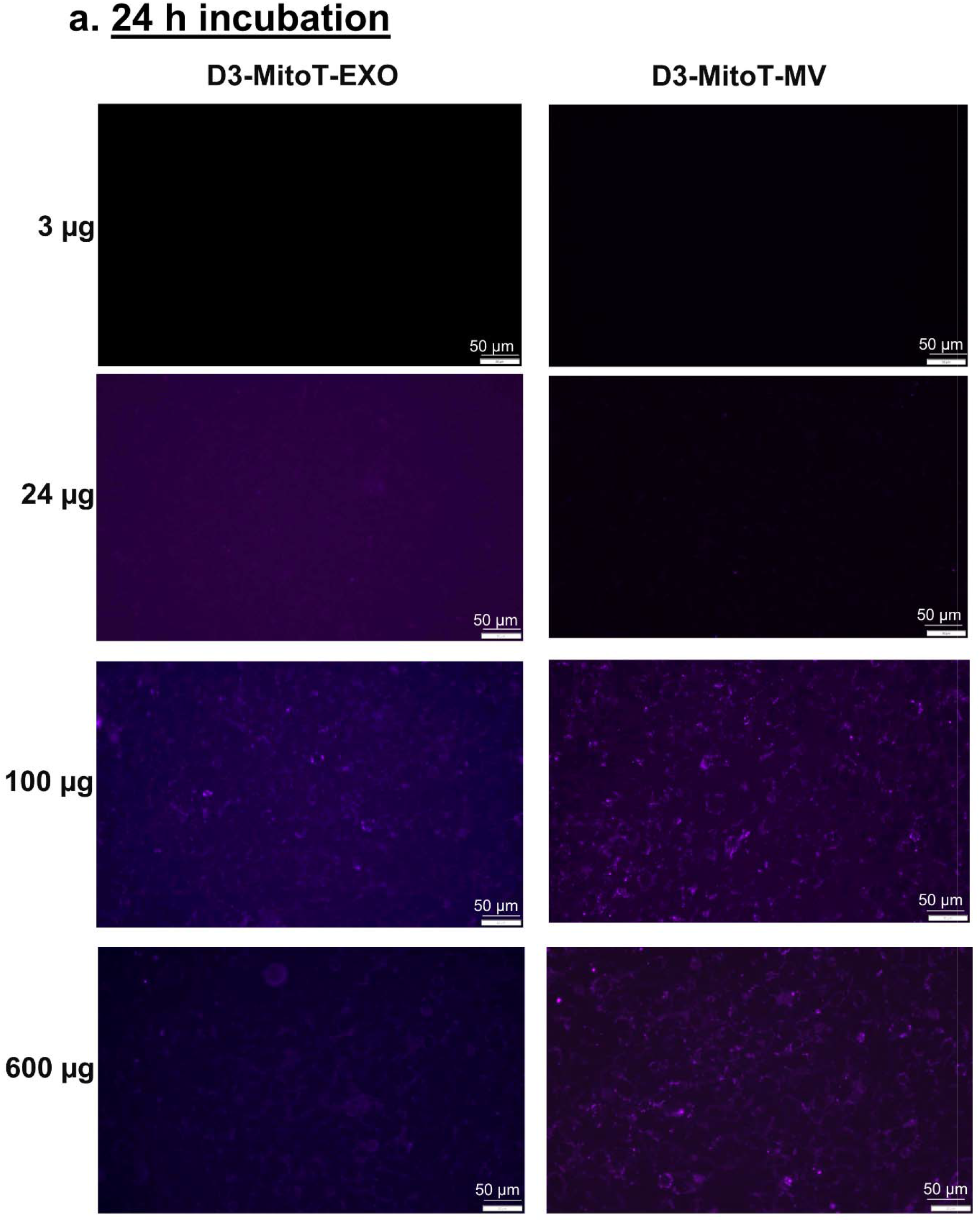
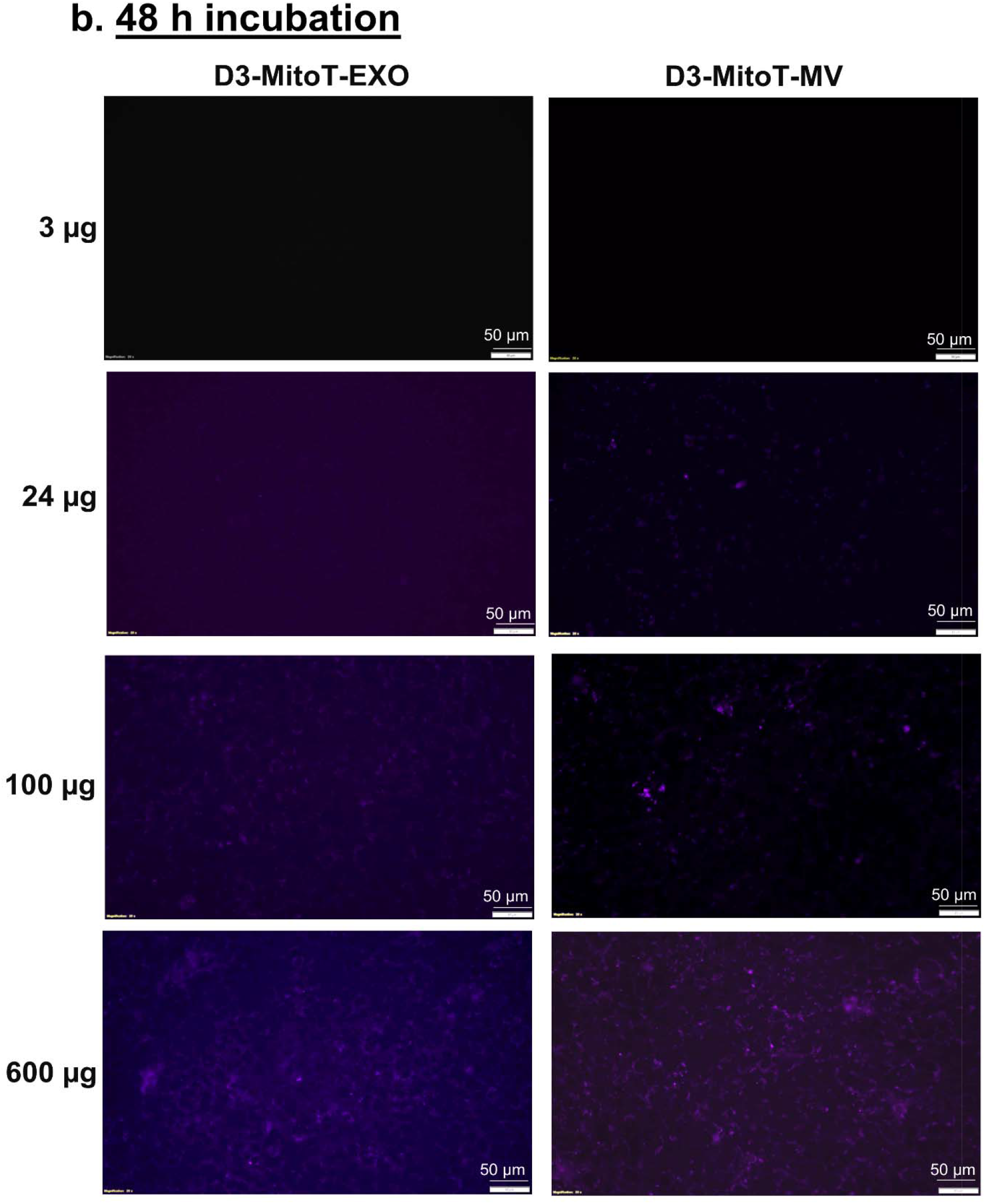
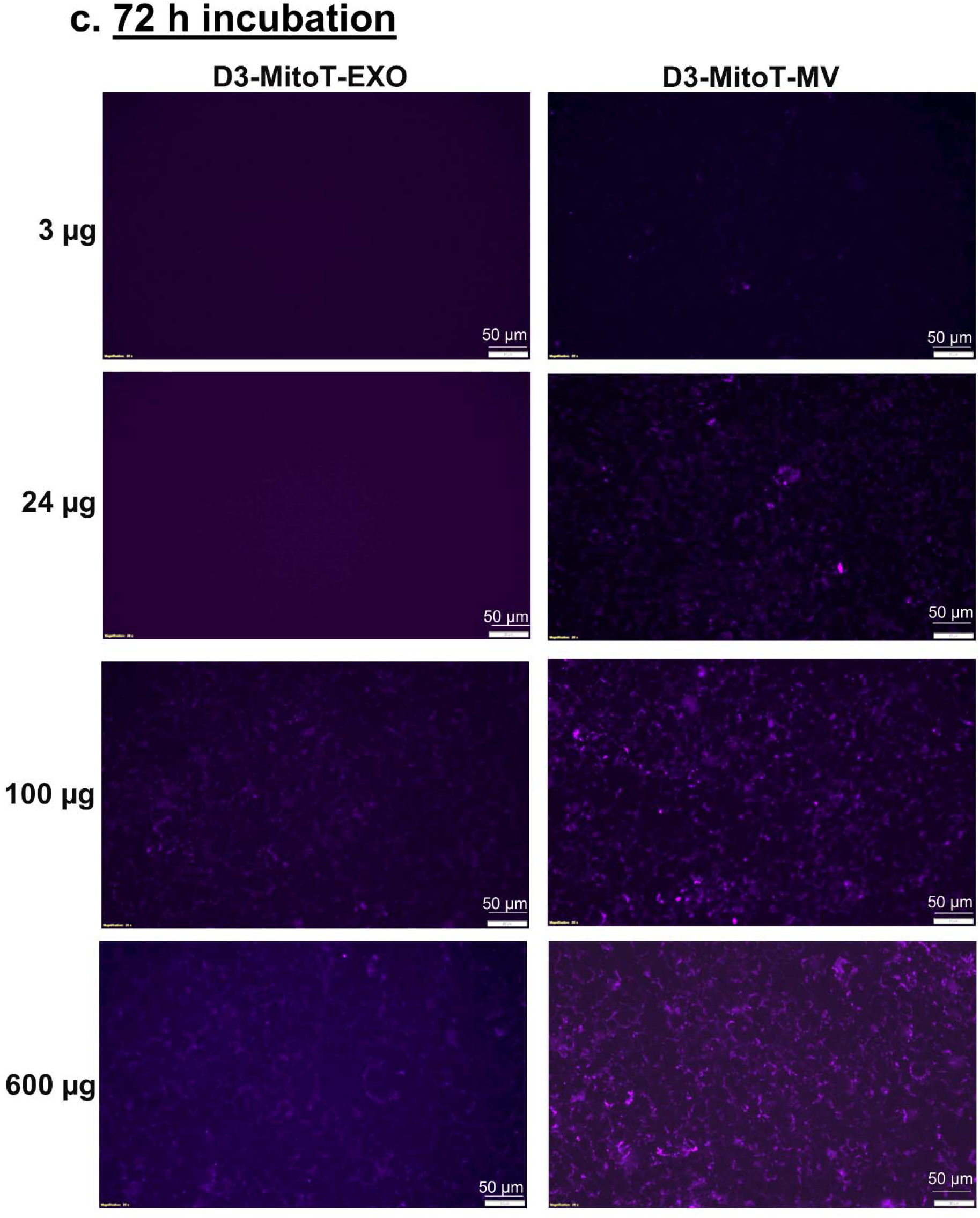
Transfer of Mitotracker-labelled mitochondria from hCMEC/D3-derived EVs to the recipient hCMEC/D3 endothelial cells. The donor/source hCMEC/D3 endothelial cell were stained with MitoTracker Deep-Red (MitoT) (250 nM for 30 min) to specifically label polarized mitochondria following which the MitoT-EVs were isolated from conditioned media. The recipient hCMEC/D3 endothelial cells were treated with D3-MitoT-EXO and D3-MitoT- MV at the indicated protein doses and observed under an Olympus IX 73 epifluorescent inverted microscope (Olympus, Pittsburgh, PA) under the Cy5 channel settings at 24 h, 48 h and 72 h post-treatment. The presented data are representative images from three independent experiments (n=4 per experiment). Scale bar = 50 μm.

**Figure 5.**
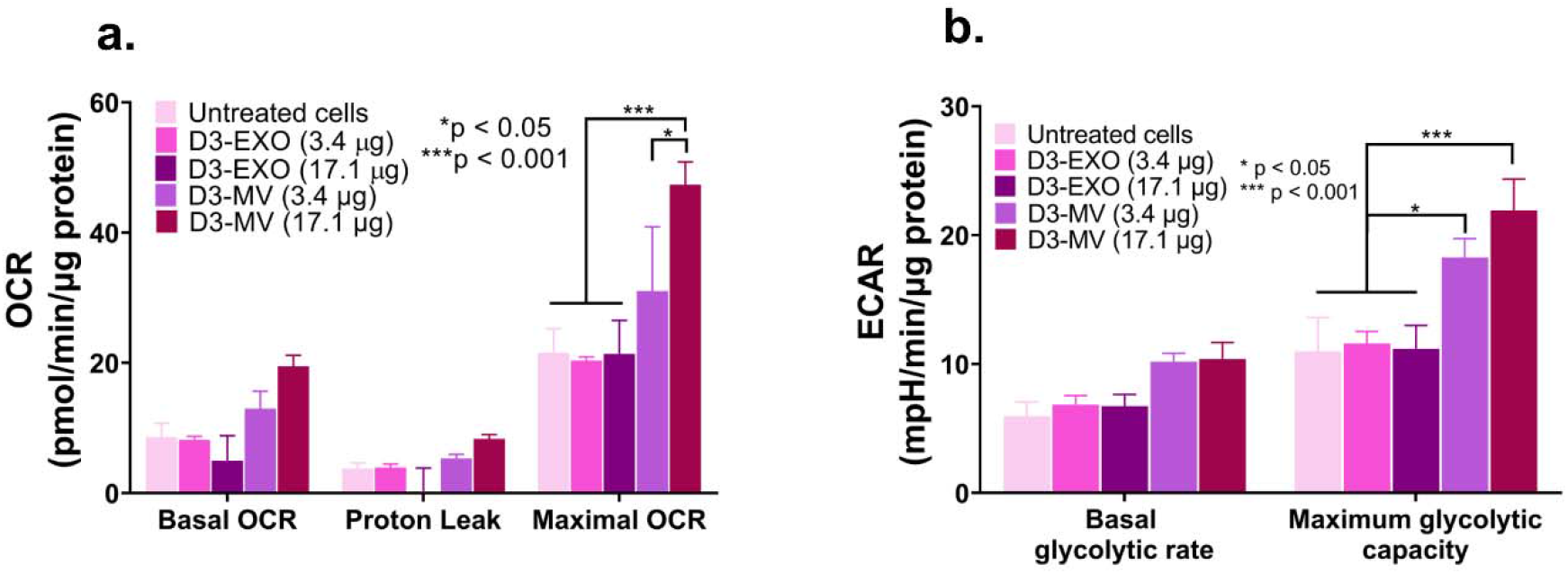
MVs increased mitochondrial function in hypoxic brain endothelial cultures. (a) Oxygen consumption and extracellular acidification rates (OCR and ECAR) were measured by treating hypoxic hCMEC/D3 cells with the indicated samples in OGD medium. We used a standard Mitochondrial Stress Test protocol to measure basal OCR followed by the addition of 2.5 μmol/L oligomycin A to measure proton leak and 0.7 μmol/L FCCP to measure maximal OCR. Basal glycolytic rate was calculated by determining the ECAR that is sensitive to 2-DG (100 mmol/L). The assay was performed in non-buffered Dulbecco’s modified Eagle medium supplemented with 25 mmol/L glucose, 1 mM pyruvate, and 2 mmol/L glutamine. All rates were normalized to cellular protein content measured using MicroBCA assay. Data are mean ± SEM, n = 3, \**p < 0.05* and \*\*\**p < 0.001* as determined using one-way ANOVA Tukey’s multiple comparisons test.

### 3.6. Mitochondrial transfer from EVs to recipient brain endothelial cells

EVs are reported to contain mitochondria, mitochondrial proteins, or mitochondrial DNA (mtDNA) and are transferable between cells [27]. Transfer of mitochondria either in the form of depolarized mitochondria that is known to be present in MVs [27] or as polarized mitochondria in EXOs [28] has been reported. Mitochondrial proteins like ATP5A were enriched in the MVs derived from brain endothelial cells [26, 84] and the umbilical cords of infants [85]. Moreover, the presence of ATP5A is also reported in the exosomal fraction isolated from murine cancer-associated fibroblasts and in serum obtained from adults with Parkinson’s disease [86, 87].

Transfer of mitochondria to the recipient cells is possible via either formation of tunnelling nanotubes, cellular fusion, GAP junctions, or microvesicles [88, 89]. F-actin microtubules or tunnelling tubes facilitate the transfer of cytoplasmic content and mitochondria to the recipient cells [90]. Mitochondrial transfer to cells via EVs can thus increase the cellular bioenergetics in the recipient cells [27, 28]. The secretion of paracrine factors, transfer of mitochondria [88, 89] and the presence of protective/antiapoptotic genetic messages or macrophage polarizing miRNAs [91, 92] are known to revive the injured cells and protect against subsequent tissue injury.

Phinney *et al*. reported that MV-mediated mitochondrial transfer under oxidative stress can improve the cell survival in the recipient macrophages cells by improving their mitochondrial bioenergetics. The authors reported that though EXOs do not contain mitochondria, they contain mtDNA that can be also transferred to the recipient cells [27]. Despite the low transfection activity of EVs containing exogenous DNA, we wanted to determine if mitochondria in EVs can be exploited for increasing the cellular energetics in the recipient endothelial cells. During ischemia/reperfusion (I/R) injury, the oxygen-glucose deprived endothelial cells lining the BBB undergo structural and functional damage leading to poor patient outcomes post-I/R injury [93]. Therefore, protection of the BBB endothelial cells is considered an effective strategy to decrease acute cell death in ischemic stroke. The presence of mitochondria organelles can increase the intracellular ATP levels and cell viability of the injured endothelial cells, thus improving mitochondrial bioenergetics, contributing to neuroprotection in an ischemic brain, and repairing brain injury [94]. The lack of ATP synthesis following oxygen and glucose deprivation sets off energy failure and loss of ionic gradients [50]. Albeit the lack of ATP cannot be equated with ischemic cell death, it is important to note that the initial trigger of ATP loss orchestrates multiple ionic, biochemical, and cellular events that lead to tissue death [95]. It is also known that glutamate excitotoxicity and calcium overload serve as additional triggers of cell death resulting in further depletion of ATP that compounds the events leading to acute cell death. Therefore, we posit that mitochondrial-transfer mediated ATP increases may serve to decrease the endothelial cell death in an *in vitro* oxygen-glucose deprivation model of stroke.

MitoTracker Deep Red (MitoT) is a carbocyanine dye that stains respiring mitochondria in live cells [96, 97] utilizing the mitochondrial membrane potential. Thus, Mitotracker Red staining is selective to polarized mitochondria and is not suggestive of depolarized or damaged mitochondria. We pre-labelled mitochondria in hCMEC/D3 and RAW 264.7 cells using MitoT and isolated EXOs and MVs (D3-MitoT-EXO, D3-MitoT-MV, RAW-MitoT-EXO, and RAW-MitoT-MV) to evaluate if the labelled mitochondria can be transferred to recipient hCMEC/D3 endothelial cells. Brain endothelial cells were treated with the indicated amounts of labelled EVs at doses ranging from 3 – 600 µg total EV protein. At a low dose of 3 μg EV protein, D3-MitoT956 EXO showed no MitoT positive signals up to 72 h whereas D3-MitoT-MV showed punctate fluorescence at 48 h of post-incubation (**Fig. 4a-c**). Furthermore, we noticed increased intracellular puncta in D3-MitoT-MV-treated cells with EV protein content ranging from 3 to 24 μg as the incubation times increased from 24 – 72 h. Cells dosed with D3-MitoT-MV from 100 μg up to 600 μg showed punctate signals at all observed time points (24-72 h). However, the signals were visible in the case of D3-MitoT-EXO only at the 100 μg protein dose at 24 h with prominent signals noted at 48 h and 72 as well. Nevertheless, there were noticeable differences in the nature of the staining observed with EXO *vs*. MV-treated cells. D3-MitoT-EXO showed faint/diffuse staining at an exposure time of 200 ms despite the higher dose of 100 μg, whereas the intracellular puncta-like staining was brighter and discrete in cells treated with D3-MitoT-MV even at a lower exposure time of 70 ms. The intensity of fluorescent signals was found to be dependent on the EV protein dose and incubation times.

We noted similar observations for RAW-MitoT-EXO and RAW-MitoT-MV at higher EV protein doses as well as at increasing incubation times from 24 to 72 h (**Fig**. **S2**). Yet another striking feature was the fact that both microvesicles, D3-MitoT-MV and RAW-MitoT-MV, showed fluorescent signals at lower treatment doses within 24 h compared to the exosomes: D3- MitoT-EXO and RAW-MitoT-EXO, that required a higher amount of protein and increased amounts of incubation time for detecting MitoT fluorescent signals. However, comparing the signal intensities, the appearance of fluorescent signals in cells was relatively earlier and more discrete with homotypic D3-MitoT-MV at lower protein contents than those incubated with the heterotypic RAW-MitoT-MV which showed fluorescent signals later and at longer exposure times. For instance, cells treated with 100 μg of D3-MitoT-MV showed higher cellular uptake than those treated with 100 μg of RAW-MitoT-MV 24-, 48-, and 72 h post-incubation. To summarize, EXOs and MVs derived from MitoT-labelled cells demonstrated accumulation of mitochondria and/or mitochondrial components in the recipient hCMEC/D3 endothelial cells. More importantly, the pattern of Mitotracker staining in the recipient cells revealed brighter puncta associated with MV transfer compared to the diffuse staining observed in the case of EXO-treated cells. The observed differences suggest that MVs are likely more efficient in transferring mitochondria compared to EXOs and aligns well with the fact that MVs incorporate mitochondria during their biogenesis. It is known that mitochondria undergo a series of dynamic changes, including biogenesis, shape changes and selective degradation and rapid transport along with cell bodies to extremities [98]. It may well be possible that these MV mitochondria are functional with their full complement of proteins, lipids, and mitochondrial DNA.

Although MV-associated mitochondria have also been previously reported [27], recent reports have demonstrated the presence of mitochondria in exosomes as well [28]. EXOs derived from airway myeloid-derived regulatory cells from both healthy and asthmatic subjects transferred mitochondria to T cells, co-localized with the mitochondrial network and regulated bioenergetics in the recipient T cells [28]. Panfoli *et al.* demonstrated that the EXOs isolated from new-born infants can produce ATP and consume oxygen with the presence of electrochemical membrane potential similar to isolated mitochondria [99]. Some mitochondrial proteins such as voltage-dependent anion channel 1 and adenosine triphosphate synthase subunit alpha were also detected in the exosomal fraction isolated from murine cancer-associated fibroblasts and in the serum obtained from adults with Parkinson’s disease isolated at 100,000*x*g and tested positive for proteins of endocytic origin [86, 87]. Some studies evaluated the entire supernatant fraction of conditioned media or plasma after removal of apoptotic bodies for MV studies [100, 101]. Zhang *et al.* used the EVs isolated from plasma deprived of cell debris and apoptotic bodies to study the mitochondrial activity. The authors found that the all fractions of EVs contained respiring mitochondria, with the highest (99.98%) being in the large-sized EVs (1 – 6 μm), intermediate (95.91%) in EVs sized 100 nm – 1 μm and low (62.72%) in small-sized EVs < 100 nm [101].

### 3.7. Measurement of mitochondrial function using Seahorse analysis

We wanted to understand if the transfer of mitochondria (**Figs. 4, S2**) affected overall mitochondrial function. We treated hypoxic endothelial cultures using EXOs or MVs and measured the basal and maximal oxygen consumption rate (OCR) as a measure of mitochondrial respiration and extracellular acidification rate (ECAR) as a measure of glycolysis. Seahorse XF Analyzer is a sensitive, robust and high-throughput technique [102] to measure mitochondrial parameters in cultured cells as well as isolated brain microvessels [43, 102–105]. Cells treated with MVs showed a significant, dose-dependent increase in the maximal OCR and ECAR compared to untreated cells and cells treated with EXOs (**Fig. 5a)**. The basal OCR was also increased by 179 and 269% in the groups treated with MVs at doses of 3.4 and 17.1 µg of MV protein (compared to untreated and EXO-treated cells), respectively. Likewise, we noted 147 and 224% increases in maximal OCR in the groups treated with MVs at a doses of 3.4 and 17.1 µg MV protein (compared to untreated and EXO-treated cells), respectively. The transfer of MV-mediated mitochondria increased the basal and maximal OCR of the hypoxic BECs (**Fig. 5a**) and as a result, the maximal glycolytic capacity also increased (**Fig. 5b**). We noted small, statistically non-significant changes in proton leak among the groups suggesting that the mitochondrial respiration is linked to ATP production via oxidative phosphorylation. An increase in the proton leak would suggest that although oxidative phosphorylation is increased, the mitochondrial function is not optimum as protons are leaking across the mitochondrial membrane.

Concomitant with the increases in OCR, the maximum glycolytic capacity of cells treated with 17.1 µg of MVs also increased by ca. 200% compared to untreated hypoxic BECs (**Fig. 5b**). It has been reported that MVs from the D3 endothelial cell line are selectively enriched with mitochondrial proteins compared to the EXO fraction [84]. Our results demonstrated that the transfer of MVs but not EXOs result in increased mitochondrial function indicated by the increased OCR and ECAR values. This MV-selective mitochondrial transfer is also supported the EV-Mitotracker images (**Fig. 4 and S2**) where the MV-treated cells showed a more discrete intracellular puncta compared to the somewhat diffuse pattern observed with the EXO-treated cells. Furthermore, our western blotting analysis (**Fig. 1g**) also demonstrated the MV were enriched with ATP5A protein compared to the EXO fraction. In summary, though published reports have reported the general trend of EV (EXO and MV)-mediated mitochondrial transfer [27, 28, 99, 101, 106–108], our data unequivocally demonstrates that the MV-mediated mitochondrial transfer increases overall mitochondrial function in a dose-dependent manner.

### 3.8. Effect of EV delivery on the ATP levels in the ischemic endothelial cells

It has been reported that depolarized mitochondria were transferred from the mesenchymal stem cells to macrophages via MVs and that these mitochondria were repurposed by undergoing fusion with the mitochondrial network of macrophages, improving their bioenergetics [27]. Thus, it is reasonable to expect that the positive effects of mitochondria-induced increases in bioenergetics would be more pronounced in ischemic cells that have impaired mitochondrial function. Therefore, we determined the effects of transferring naïve EVs in brain endothelial cells subjected to an ischemic attack using an *in vitro* oxygen-glucose deprivation (OGD) model of stroke. We used a CellTiter Glo-Luminescent Cell Viability assay (referred to as “ATP assay” henceforth) to measure the resulting ATP levels upon EV exposure. We chose the ATP assay to determine the effects of EV exposure as it is a rapid and sensitive technique for evaluating the cell viability of the treated cells [109, 110] and the ATP readout is directly proportional to the number of cells in culture [111]. The cytoplasmic mitochondrial volume of rat brain endothelial cells was almost two to four-fold higher compared to the other non-brain endothelial cells (2 to 5%), suggesting that the brain endothelial cells have a higher metabolic activity [48]. Additionally, hypoxic conditions lead to disruption of tight junctions and apoptosis in BECs further increasing the need for mitochondrial metabolism for endothelial survival [49]. Oxygen-glucose deprivation reduces oxidative phosphorylation and induces energy failure [27]. Recovery of bioenergetics in cells is indicated by their ability to generate mitochondrial ATP in coupled with proton leak and/or generation of reactive oxygen species [27]. We first optimized the exposure time of cells to determine the reproducibility of simulating OGD-induced cell death in the hCMEC/D3 brain endothelial cells. We determined that a 4 h OGD exposure was sufficient to mediate at least 50% cell death as measured using the ATP assay (**Fig. S3**).

We first exposed healthy hCMEC/D3 endothelial cells (cultured under normoxic conditions) to different EV protein doses ranging from 14.6 to 174.5 μg (per 0.32 cm2/well in a 96-well plate). These selected doses were equivalent to 50, 100, 200, 400 and 600 μg per 0.85 cm2 area of 48-well plate that was previously used in the MitoT-EV study (**Fig. 4 and S2**). Seventy-two hours post-incubation with normoxic monolayers, no significant differences were found in the ATP levels (and the resulting cell viabilities) in cells that were treated with both D3- and RAW-EXOs and D3-MV groups at 14.6 and 29.1 μg and compared to the untreated cells (**Fig. 6a**). The cell viabilities however decreased with an increase in the treatment dose of EXO dose from 58.2 to 174.5 μg and the case of D3-MVs, from 29.1 μg and upward. A previous study reported no cytotoxicity of milk-derived EVs up to doses of 200 μg protein/mL in Caco-2 intestinal monolayers since these cells naturally absorb digestive products [112]. Hansen *et al*. observed decreased Caco-2 cell viability when treated with 50 μg/mL of Alexa Fluor-labelled EVs isolated from milk after 6 h but the cell viability was regained after 24 h, which however were not confirmed in the successive experiments [113].

**Figure 6.**
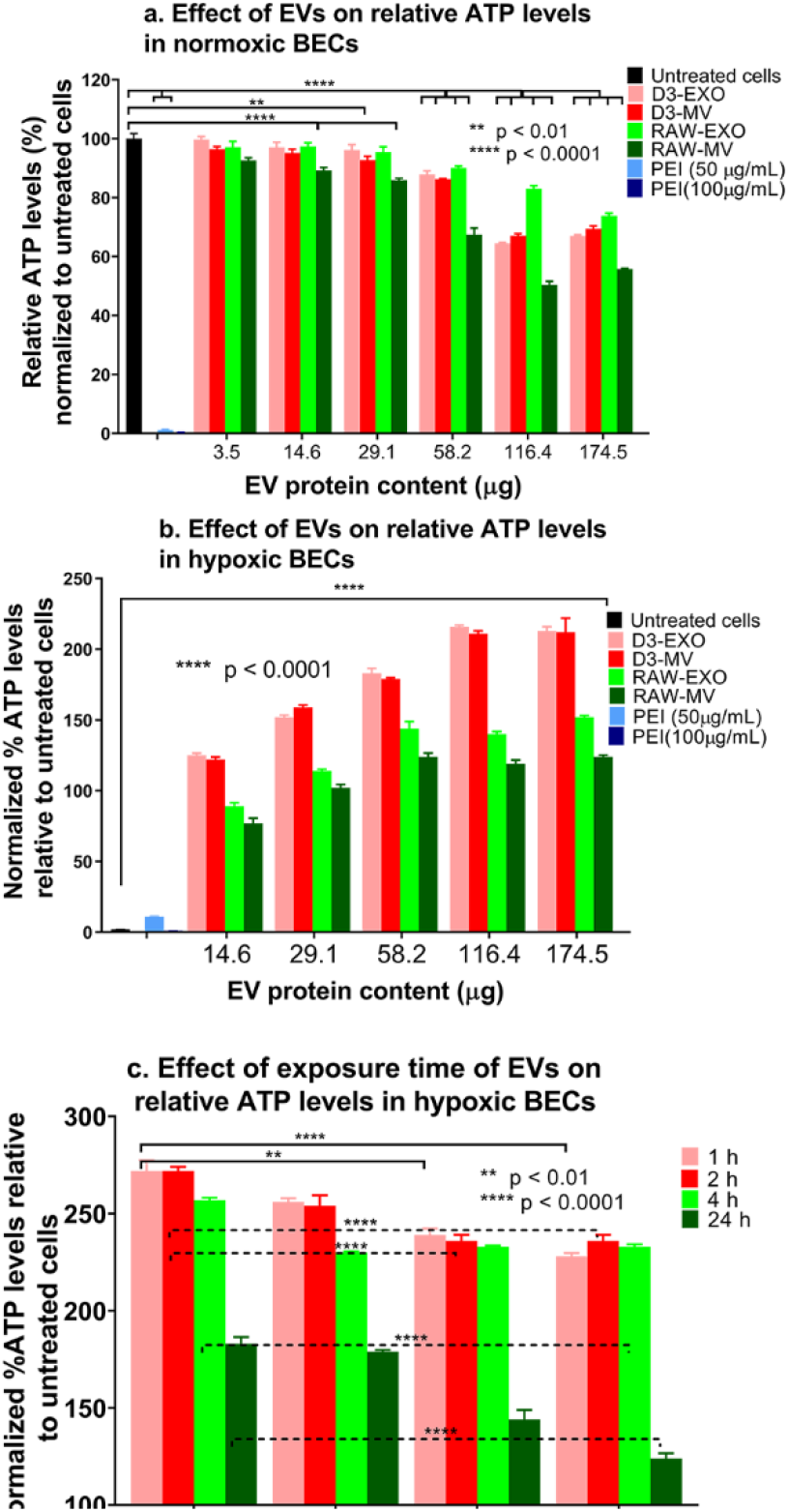
Effects of EV exposure on the ATP levels of hCMEC/D3 endothelial cells under normoxic and hypoxic (OGD) conditions. (**a**) Normoxic hCMEC/D3 endothelial cells were treated with the indicated EV protein content for 72 h. (**b**) hCMEC/D3 endothelial cells were subjected to 4 h of OGD by exposing the cells in a sealed hypoxia chamber (90% N_2_, 5% H_2_, 5% CO_2_) and glucose-free media at 37 °C in a humidified incubator. OGD exposed hCMEC/D3 endothelial cells were treated with the indicated amounts of naïve D3-EVs and RAW-EVs for 24 h under 21% O_2_ in a humidified incubator (normoxic conditions). Untreated OGD cells were cultured in glucose-free media under normoxic conditions. (**c**) Effect of exposure time on the resulting ATP levels in hypoxic hCMEC/D3 endothelial cells. Hypoxic monolayers were treated with the indicated amounts of EVs for the indicated periods. Untreated OGD cells were cultured in glucose-free media 4 h post-OGD under normoxic conditions. Unless indicated otherwise, normoxic cells treated with polyethyleneimine (PEI) for 4 h were considered as a positive control. In all cases, the effects of treatment were determined using an ATP assay. Data are represented as mean_±_SD (n = 4). Statistical comparisons to the normoxic/hypoxic groups were made using one-way ANOVA Bonferroni’s multiple comparisons test.

All types of EVs regardless of parent cell source resulted in increased cellular ATP levels relative to control, untreated hypoxic cells under OGD conditions (**Fig. 6b**). The increased relative ATP levels were likely indicative of greater cell survival at all tested EVs doses. The ATP levels were significantly higher in recipient cells when incubated with all doses of the different EVs compared to the control group. Incubation of OGD-exposed endothelial cells with D3- and RAW-EVs increased the ATP levels by nearly 100 to 200-fold as a function of the EV protein dose. At the same time, no differences were found when the recipient cells were treated with D3-EXO and D3-MV. However, the ATP levels of D3-EVs post-incubation were higher than the ATP levels of RAW-EVs post-incubation (**Fig. 6b and 6c**). Our results showed that at least 14.6 μg of total EV protein is required to increase the cellular ATP levels. A similar observation was noted when EVs derived from human Wharton’s jelly mesenchymal stem cells were used to treat OGD-exposed mouse neuroblastoma cells [114]. Another study [115] reported that EV doses of 200 μg yielded the maximum cell proliferation, while 50 μg of EVs was the minimal effective dose to increase cell proliferation in neural stem cells under OGD conditions.

Our results further confirmed that EVs can increase cellular ATP levels in a dose-dependent manner. A modest increase in ATP levels was observed at the highest protein dose of 116.4 μg with D3-derived EVs and at doses greater than 58.2 μg with RAW-derived EVs. It is likely that at the higher doses, the cells may have shown a reduced uptake with time causing the observed plateau effects. A saturation in uptake or intake equivalent to intracellular processing of EVs has been reported after 18 h [113] especially in the case of milk-derived EVs [116]. Saari *et al*. had similar observations with paclitaxel-loaded EVs which showed increased toxicity with an increase in concentrations in LNCaP and PC-3 cell lines and their respective EVs. However, after incubating cells with 10^9^ paclitaxel loaded-EVs/mL, a saturation point with maximum cytotoxic effect was reached at 24 h [117]. Meanwhile, HEK293T-derived EVs showed early uptake within 2 h with a peak at 12 h and then decreased up to 24 h. The lower values may also be due to the exocytosis of EVs after 24 h [116].

The protective effects of the EVs appeared to reach their maximum at about 2 h post-incubation (**Fig. 6c**) in hypoxic cultures. As described in the above paragraph, EV uptake also shows a saturation and a time-dependent uptake. Hansen *et al.* observed a plateau in EV uptake 18 h post-exposure [113]. The saturation effects are likely due to the active, energy-dependent endocytic uptake of EVs [118–120] and not via passive membrane fusion, which caused an inhibition in the uptake of EVs [121]. Noteworthy, the resulting ATP levels were consistently higher with D3-derived EVs compared to the RAW-derived EVs suggesting that under cell duress, the recipient D3 brain endothelial cells preferentially internalized the homotypic D3- derived EVs as opposed to the heterotypic RAW macrophage-derived EVs (**Fig. 6b**).

ATP is the most common intracellular energy source and importantly, in high energy consuming-tissues like the heart and brain, mitochondria produce 80-90% of the cell’s ATP [122]. Depletion of ATP levels is thought to prevent efficient post-ischemic repair [122]. The EV-mediated ATP increases suggest that this may be a promising approach to decrease acute cell death and activate ischemic repair pathways to limit post-stroke damage. Mitochondrial transfer helps to rescue metabolism and protects the neurons and other brain cells from tissue injury. Brain endothelial cells also take up extracellular mitochondria derived from endothelial progenitor cells under OGD conditions [123]. EXOs are also reported to contain mitochondria or mtDNA which can be transferred across distant cells [27, 28, 99, 106–108]. Mitochondrial DNA and intracellular ATP were upregulated in the oxygen-glucose deprived endothelial progenitor cells when treated with endothelial progenitor cell-derived mitochondria [123]. Mitochondrial DNA can also be imported into mitochondria irrespective of the mitochondrial membrane potential [124]. The mammalian mtDNA is essential to couple respiration to ATP synthesis and oxidative photophosphorylation [125]. It also encodes functional RNAs in intramitochondrial translation.

Transfer of mtDNA can be of crucial significance in increasing the mitochondrial load in the recipient cells. The mtDNA can be imported into the mitochondria once it reaches the cytosol of the recipient cell [106]. EVs derived from cardiomyocytes containing chromosomal DNA were transferred to the nuclei or cytosol of the recipient fibroblasts [126]. Transfer of intact/full mitochondria can also import mtDNA and provide a template for synthesis of DNA and RNA [124]. Adipose stem cell-derived EXOs reverted mitochondrial dysfunction by contributing complex 1 to the electron transfer system and coupling efficiency as well as by restoring mitochondrial membrane potential [127]. Mesenchymal stem cell-derived EXO containing glycolytic enzymes restored the glycolysis and ATP levels with reduced oxidative stress in mice subjected to 30 min of ischemia followed by reperfusion [128] and also increased the levels of extracellular ATP [129]. Our data and these findings thus suggest that EVs-derived from D3 endothelial and RAW cells may increase the cellular bioenergetics of brain endothelial cells in an animal model of ischemic stroke through mitochondrial transfer, providing an extracellular source of glycolytic enzymes and/or the transfer of mtDNA.

### 3.9. Uptake of MitoT-EVs by cortical and granule neurons in mice acute brain cortical and hippocampal slices

It is known that both brain endothelial and neuronal cultures equally and rapidly respond to ischemic injury [130, 131] and therefore, we wanted to understand the effects of EVs on neurons (**Figure 7**). Injury to a specific region of the brain microvasculature (composed of endothelial cells lining the BBB) can also damage adjacent neurons [130]. We used acute brain slices as an *ex vivo* model to determine if the EVs show uptake into the neurons in cortical and hippocampal slices. Acute brain slices preserve the natural cytoarchitecture and are a robust model to study the detailed cellular, molecular and circuitry level analysis of neuronal function [132]. Cortical slices obtained from non-surgical control mice were incubated in D3-MitoT-EVs for 2 h at 37 °C. In slices incubated with D3-MitoT-MV, we observed a punctate intracellular staining within neurons and vascular staining in both the cortical (**Fig. 7a**) and hippocampal (dentate gyrus, **Fig. 7b**) slices. Analysis of mean intensity normalized to average control, untreated slices show an increase in D3-MitoT-MV intensity in the cortex **(Fig. 7c**), increasing from 1.1±0.2 in control slices to 2.5±0.6 in MV treated slices (p<0.1, n=4). No appreciable increase was detected in D3-MitoT-EXO treated hippocampal slices, 1.3±0.3 (n=4). In the D3-MitoT-EXO and control conditions, we found similar levels of nonspecific punctate staining in both brain regions (**Fig. 7a, 7b**). Similarly, analysis of mean intensity normalized to average control levels show an increase in D3-MitoT-MV intensity in the hippocampus (**Fig. 7d**), increasing from 1.0±0.2 in control slices to 1.7±0.4 in MV-treated slices (p<0.1, n=4). No appreciable increase was detected in D3-MitoT-EXO treated hippocampal slices, 1.5±0.7 (n=4). These results suggest that MVs but not EXOs transfer polarized mitochondria to neuronal and endothelial cells in cortical and hippocampal brain slices. These results align well with the MV-mediated mitochondrial transfer into endothelial cultures (**Figures 4 and S2**) and the selective MV-mediated increases in mitochondrial function (**Figure 5**).

**Figure 7.**
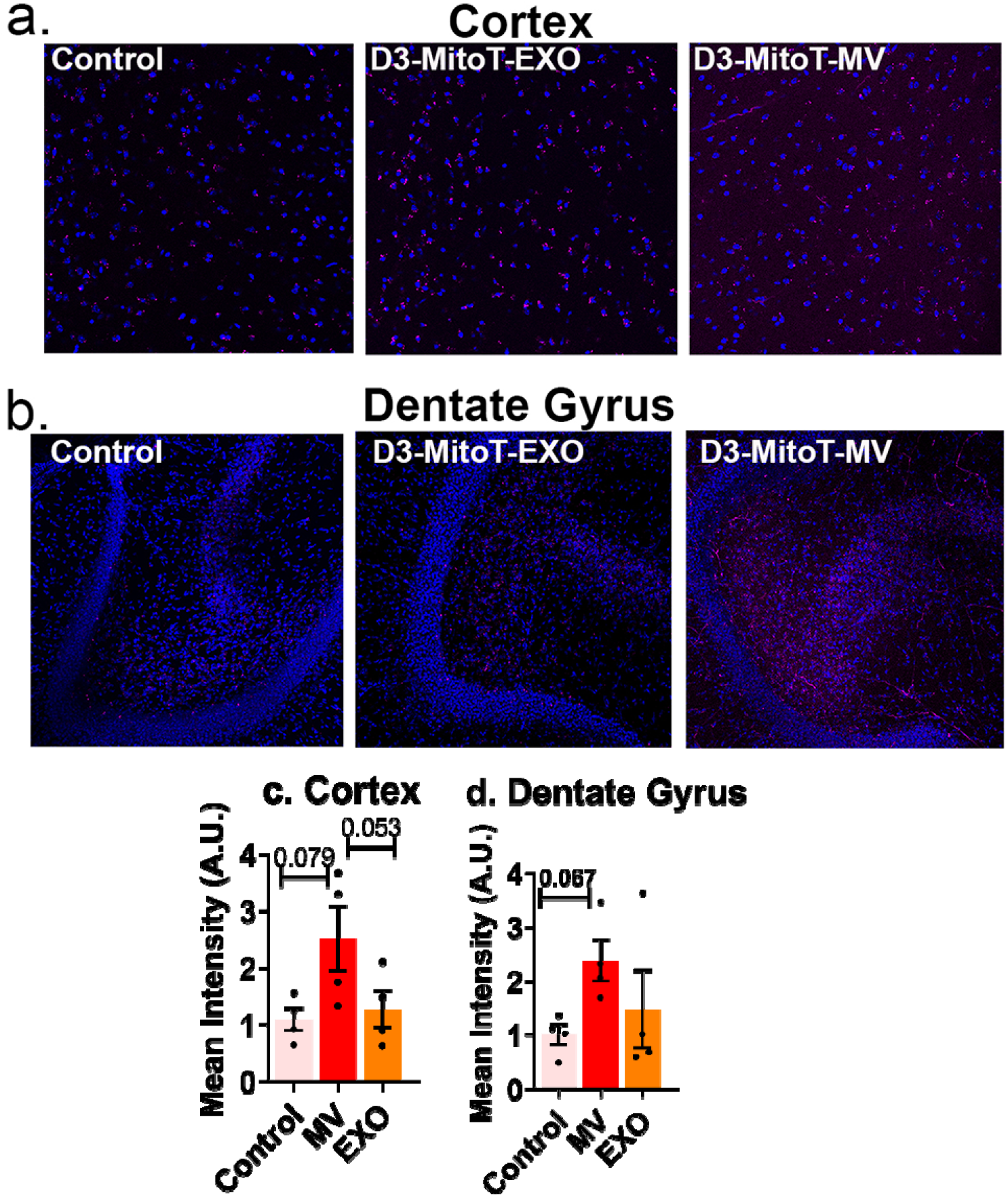
Uptake of MitoT-EVs by acute brain slices. Acute cortical and hippocampal slices from mice subjected to sham middle cerebral artery occlusion procedure were left either untreated (control) or incubated in 50 µg/mL of D3-MitoT-EXO or D3-MitoT-MV for 2 h at 37 °C. Slices were fixed, counterstained with Hoechst 33258 (blue), and visualized on a confocal microscope. Intracellular punctate staining within neurons (magenta) and vascular staining were evident in slices from the cortex (*a*) and dentate gyrus region of the hippocampus (*b*) in the D3-MitoT-MV treated condition. The control and D3-MitoT-EXO conditions exhibited similar levels of nonspecific staining in both regions (*a and b*). Mean intensity values were normalized to control slices and statistical analysis was done using GraphPad Prism 9.1.2 software (*c and d*).

Although these images were obtained from healthy mice, the potential of using EV and MVs to increase cellular energy levels in injured neurons is intriguing and has significant implications in treating ischemic injury *in vivo*. Neurons can be injured due to oxidative stress and inflammation as a result of cell death following ischemia reperfusion injury. A recent report demonstrated that mitochondrial transfer from mesenchymal stem cells co-cultured with mice primary neurons rescued the H_2_O_2_-injured neurons and improved metabolism [133]. Seahorse analysis revealed that the mitochondrial respiratory parameters such as basal respiratory rate, spare respiratory capacity, ATP production and proton leak in the injured neurons were significantly improved upon mitochondrial transfer [133]. Impaired and depolarized mitochondria result in decreased ATP, increased reactive oxygen species and calcium overloading which opens up the membrane permeability transition pore releasing cytochrome C and eventually leads to apoptosis. Mitochondrial fusion with the injured cells causes rapid exchange of mitochondrial DNA, mitochondrial membranes and mitochondrial metabolites within mitochondrial network and repair the damaged ones. Thus, mitochondrial transfer increases the chances of neuronal cell survival upon ischemic attack.

### 3.10. Delivery of ATP5A protein via EV/ATP5A complexes to hypoxic BECs

It is known that D3-derived microvesicles were highly enriched in ATP5A1 [26, 84], a subunit of the mitochondrial ATP synthase protein that catalyzes ATP synthesis [134]. From an engineering perspective, we tested if we could load exogenous ATP5A into EVs to further buttress its capability to deliver ATP5A protein. ATP5A1 is a mitochondrial [135], nucleus-encoded protein [136] biosynthesized by the mitochondria to produce ATP from ADP in presence of a proton gradient. Reduced supply of blood and oxygen during ischemia causes an imbalance of the energy production and depletes the high energy phosphates, ATP. Direct infusion of exogenous ATP is not possible due to its anionic charge and hydrophilic nature which forbids delivery through the cellular membrane. Moreover, the systemic half-life of ATP is short (< 40s) due to degradation by ectonucleotidases [137]. Hypoxia/reperfusion of cells decreases the ATP5A/ mitochondrial-encoded protein cytochrome-c oxidase I ratio [136] and ATP5A mRNA levels [138] further exacerbating ischemic injury.

ATP5A is a large cationic protein (MW 59.8 kD, pI 9.2) that cannot diffuse or penetrate through the cell membrane. ATP5A contains 68 cationic residues (arginine and lysine) at a physiological pH of 7.4 that can form a complex with the negatively-charged EV membranes via electrostatic interactions. The formation of the EV/ATP5A complexes was confirmed using a native polyacrylamide gel (**Fig. 8a**). Coomassie dye stained both the ATP5A and EV proteins. Under native PAGE conditions, ATP5A is slightly positively charged at the running buffer pH of 8.3. On the other hand, the negatively-charged EVs ranging with a surface zeta potential of -4 and -12 mV (**Fig. 8a**) will migrate towards the anode. While the cationic free ATP5A protein stayed at the loading point, EV protein: ATP5A complexes at a weight/weight ratio (w/w) of 5:1 resulted in the retention of the formed complexes at the loading point while the excess free EVs slightly migrated towards the anode (**+**) confirming the neutralization of protein charges by EV membranes. It should be noted that this pattern of migration of EV/protein complexes was consistent with previously reported findings on EV complexes with a similar cationic protein, brain-derived neurotrophic factor [21].

**Figure 8.**
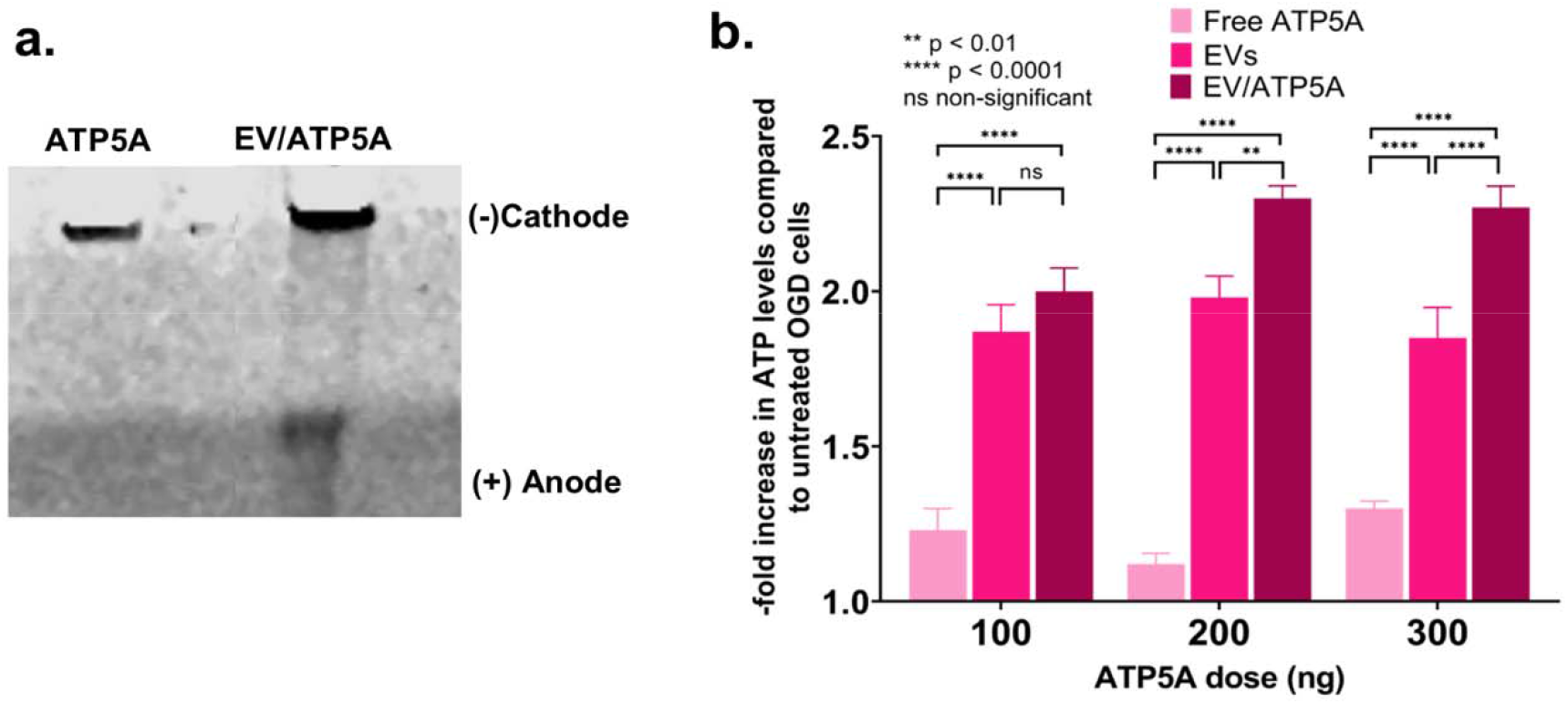
Formation of EV/ATP5A complexes confirmed using native gel electrophoresis. (*a*) Native PAGE analysis of the EV/ATP5A complexes. 0.5 μg of the indicated samples were electrophoresed in a 4-10% of native PAGE gel and the bands were visualized using Bio-safe Coomassie dye. (*b*) Confluent hCMEC/D3 cells (16,000/well) in 96-well plates were exposed to OGD conditions for 4 h following which the media was replaced with 100 μL of the indicated samples. Cells were incubated for 24 h and washed once in 1*x* PBS prior to measuring cellular ATP levels using a Cell Glo luminescence assay. Data represent average ± SD (n=4).

The effect of EV/ATP5A complexes on the resulting ATP levels in OGD-exposed hCMEC/D3 is shown in **Fig. 8b**. Incubation of free ATP5A1 with endothelial cells exposed to OGD increased the cellular ATP levels. However, increasing the dose of free ATP5A1 from 100 to 300 ng did not result in a significant increase in the ATP levels that almost remained constant regardless of the ATP5A dose. As observed and discussed in section *3.8*., naïve EVs increased the ATP levels of the endothelial cells exposed to OGD. However, the increase in the ATP levels is not significant with an increase in the naïve EV doses (500, 1000 and 1500 ng) added in amounts equivalent to those in EV/ATP5A complexes. The EV/ATP5A complexes containing 100 ng of ATP5A also did not result in a significant increase in the ATP levels compared to cells treated with naïve EVs. However, the ATP levels increased significantly in cells treated with EV/ATP5A complexes containing 200 ng of ATP5A protein. A further increase in the ATP5A dose to 300 ng in the EV/ATP5A complexes showed elevated ATP levels. Our observations point at a trend that higher ATP5A protein doses (200 and 300 ng) delivered via EVs increased ATP levels by ca. 2.3-fold compared to about 1.9-fold in the case of naïve EVs.

## 4. Conclusions

We suggest that the natural overlap of exosomal biogenesis and the intracellular trafficking pathways of DNA-EVs may explain the greater DNA loading in smaller EXOs compared to the larger MVs, in contrast to previously reported findings. Gene ontology and pathway enrichment analyses revealed that EVs overexpressed glycolytic genes and pathways. We have demonstrated, for the first time, that homotypic, endothelial-derived EVs, result in a greater extent of mitochondrial transfer to the recipient’s brain endothelial cells and resulting ATP increases, compared to heterotypic, macrophage-derived EVs. Transfer of microvesicles, but not exosomes, resulted in increased mitochondrial functions in the recipient cells. We have also demonstrated microvesicle-mediated mitochondrial transfer to neurons in acute brain cortical and hippocampal slices. Our findings suggest that EV carriers have immense potential to increase cellular- and mitochondrial bioenergetics in the endothelial cells lining the BBB and neurons. Therefore, EVs are attractive drugs for the treatment of not only ischemic stroke but also for treating brain disorders wherein the permeability of BBB is altered.

## Acknowledgments

The authors are grateful to Dr. Rehana Leak (DU) for the inspiring discussions. The authors are thankful to Mr. Tarun Bhatia in the Leak Lab for technical assistance and Dr. Jelena Janjic for allowing the use of Malvern Zetasizer Nano. The authors are also thankful to the Biological Sciences Department of Duquesne University for Equipment Support and the Biomedical Mass Spectrometry Center, University of Pittsburgh School of Medicine for the proteomics study.

## Appendix A. Supplementary data

Supplementary data related to this article can be found in the accompanying Word file.

