## Supplementary Information for "Microvesicles Transfer Mitochondria and Increase Mitochondrial Function in Brain Endothelial Cells"

**Table S1**. Top 100 proteins identified in the isolated EVs compared to Vesiclepedia database

*Top 100 EV proteins listed in the Vesiclepedia database were downloaded on 02/14/2021.*

| **Vesiclepedia information** | | **Expression in** | |
| --- | --- | --- | --- |
| **No.** | **Gene Symbol** | **Naïve D3-EV** | **Naïve RAW-EV** |
| 1 | **PDCD6IP** | Not detected | Q80Y09_MOUSE Pdcd6ip protein OS=Mus musculus OX=10090 GN=Pdcd6ip PE=2 SV=1 |
| 2 | **GAPDH** | G3P_HUMAN Glyceraldehyde-3-phosphate dehydrogenase OS=Homo sapiens OX=9606 GN=GAPDH PE=1 SV=3 | D2KHZ9_MOUSE Glyceraldehyde-3-phosphate dehydrogenase OS=Mus musculus OX=10090 GN=GAPDH PE=2 SV=1 |
| 3 | **HSPA8** | Not detected | Q3UBA6_MOUSE Uncharacterized protein OS=Mus musculus OX=10090 GN=Hspa8 PE=2 SV=1 |
| 4 | **ACTB** | Not detected | B2RRX1_MOUSE Actin, beta OS=Mus musculus OX=10090 GN=Actb PE=2 SV=1 |
| 5 | **ANXA2** | ANXA2_HUMAN Annexin A2 OS=Homo sapiens OX=9606 GN=ANXA2 PE=1 SV=2 | Not detected |
| 6 | **CD9** | Not detected | CD9_MOUSE CD9 antigen OS=Mus musculus OX=10090 GN=Cd9 PE=1 SV=2 |
| 7 | **PKM** | KPYM_HUMAN Pyruvate kinase PKM OS=Homo sapiens OX=9606 GN=PKM PE=1 SV=4 | KPYM_MOUSE Pyruvate kinase PKM OS=Mus musculus OX=10090 GN=Pkm PE=1 SV=4 |
| 8 | **HSP90AA1** | HS90A_HUMAN Isoform 2 of Heat shock protein HSP 90-alpha OS=Homo sapiens OX=9606 GN=HSP90AA1 | Not detected |
| 9 | **ENO1** | ENOA_HUMAN Alpha-enolase OS=Homo sapiens OX=9606 GN=ENO1 PE=1 SV=2 | ENOA_MOUSE Alpha-enolase OS=Mus musculus OX=10090 GN=Eno1 PE=1 SV=3 |
| 10 | **ANXA5** | Not detected | ANXA5_MOUSE Annexin A5 OS=Mus musculus OX=10090 GN=Anxa5 PE=1 SV=1 |
| 11 | **HSP90AB1** | Not detected | 1) ENPL_MOUSE Endoplasmin OS=Mus musculus OX=10090 GN=Hsp90b1 PE=1 SV=2  2) HS90B_MOUSE Heat shock protein HSP 90-beta OS=Mus musculus OX=10090 GN=Hsp90ab1 PE=1 SV=3 |
| 12 | **CD63** | Not detected | Not detected |
| 13 | **YWHAZ** | 1433Z_HUMAN 14-3-3 protein zeta/delta OS=Homo sapiens OX=9606 GN=YWHAZ PE=1 SV=1 | Not detected |
| 14 | **YWHAE** | 1433E_HUMAN 14-3-3 protein epsilon OS=Homo sapiens OX=9606 GN=YWHAE PE=1 SV=1 | Not detected |
| 15 | **EEF1A1** | Not detected | EF1A1_MOUSE Elongation factor 1-alpha 1 OS=Mus musculus OX=10090 GN=Eef1a1 PE=1 SV=3 |
| 16 | **PGK1** | Not detected | PGK1_MOUSE Phosphoglycerate kinase 1 OS=Mus musculus OX=10090 GN=Pgk1 PE=1 SV=4 |
| 17 | **CLTC** | CLH1_HUMAN Clathrin heavy chain 1 OS=Homo sapiens OX=9606 GN=CLTC PE=1 SV=5 | Q5SXR6_MOUSE Clathrin heavy chain OS=Mus musculus OX=10090 GN=Cltc PE=1 SV=1 |
| 18 | **PPIA** | Not detected | PPIA_MOUSE Peptidyl-prolyl cis-trans isomerase A OS=Mus musculus OX=10090 GN=Ppia PE=1 SV=2 |
| 19 | **SDCBP** | Not detected | Not detected |
| 20 | **ALDOA** | ALDOA_HUMAN Fructose-bisphosphate aldolase A OS=Homo sapiens OX=9606 GN=ALDOA PE=1 SV=2 | A6ZI44_MOUSE Fructose-bisphosphate aldolase OS=Mus musculus OX=10090 GN=Aldoa PE=1 SV=1 |
| 21 | **EEF2** | Not detected | EF2_MOUSE Elongation factor 2 OS=Mus musculus OX=10090 GN=Eef2 PE=1 SV=2 |
| 22 | **ALB** | ALBU_HUMAN Serum albumin OS=Homo sapiens OX=9606 GN=ALB PE=1 SV=2 | ALBU_MOUSE Serum albumin OS=Mus musculus OX=10090 GN=Alb PE=1 SV=3 |
| 23 | **TPI1** | TPIS_HUMAN Isoform 2 of Triosephosphate isomerase OS=Homo sapiens OX=9606 GN=TPI1 | TPIS_MOUSE Triosephosphate isomerase OS=Mus musculus OX=10090 GN=Tpi1 PE=1 SV=4 |
| 24 | **VCP** | Not detected | TERA_MOUSE Transitional endoplasmic reticulum ATPase OS=Mus musculus OX=10090 GN=Vcp PE=1 SV=4 |
| 25 | **CFL1** | Not detected | COF1_MOUSE Cofilin-1 OS=Mus musculus OX=10090 GN=Cfl1 PE=1 SV=3 |
| 26 | **MSN** | Not detected | MOES_MOUSE Moesin OS=Mus musculus OX=10090 GN=Msn PE=1 SV=3 |
| 27 | **ATP1A1** | Not detected | AT1A1_MOUSE Sodium/potassium-transporting ATPase subunit alpha-1 OS=Mus musculus OX=10090 GN=Atp1a1 PE=1 SV=1 |
| 28 | **PRDX1** | Not detected | Not detected |
| 29 | **MYH9** | MYH9_HUMAN Myosin-9 OS=Homo sapiens OX=9606 GN=MYH9 PE=1 SV=4 | MYH9_MOUSE Myosin-9 OS=Mus musculus OX=10090 GN=Myh9 PE=1 SV=4 |
| 30 | **EZR** | Not detected | Not detected |
| 31 | **CD81** | Not detected | Not detected |
| 32 | **ANXA6** | Not detected | Not detected |
| 33 | **FLOT1** | Not detected | Not detected |
| 34 | **YWHAB** | Not detected | Not detected |
| 35 | **LDHB** | Not detected | Not detected |
| 36 | **SLC3A2** | 4F2_HUMAN 4F2 cell-surface antigen heavy chain OS=Homo sapiens OX=9606 GN=SLC3A2 PE=1 SV=3 | 4F2_MOUSE 4F2 cell-surface antigen heavy chain OS=Mus musculus OX=10090 GN=Slc3a2 PE=1 SV=1 |
| 37 | **GNB1** | Not detected |  |
| 38 | **PFN1** | Not detected | PROF1_MOUSE Profilin-1 OS=Mus musculus OX=10090 GN=Pfn1 PE=1 SV=2 |
| 39 | **TSG101** | Not detected | Not detected |
| 40 | **YWHAQ** | 1433T_HUMAN 14-3-3 protein theta OS=Homo sapiens OX=9606 GN=YWHAQ PE=1 SV=1 | Not detected |
| 41 | **GNAI2** | Not detected | Not detected |
| 42 | **CLIC1** | Not detected | Q542F1_MOUSE Chloride intracellular channel protein OS=Mus musculus OX=10090 GN=Clic1 PE=1 SV=1 |
| 43 | **ANXA1** | ANXA1_HUMAN Annexin A1 OS=Homo sapiens OX=9606 GN=ANXA1 PE=1 SV=2 | Not detected |
| 44 | **ITGB1** | Not detected | 1) ITB2_MOUSE Integrin beta-2 OS=Mus musculus OX=10090 GN=Itgb2 PE=1 SV=2  2) ITB1_MOUSE Integrin beta-1 OS=Mus musculus OX=10090 GN=Itgb1 PE=1 SV=1 |
| 45 | **LDHA** | LDHA_HUMAN Isoform 3 of L-lactate dehydrogenase A chain OS=Homo sapiens OX=9606 GN=LDHA | A0A1B0GSR9_MOUSE L-lactate dehydrogenase OS=Mus musculus OX=10090 GN=Ldha PE=1 SV=1 |
| 46 | **FASN** | FAS_HUMAN Fatty acid synthase OS=Homo sapiens OX=9606 GN=FASN PE=1 SV=3 | FAS_MOUSE Fatty acid synthase OS=Mus musculus OX=10090 GN=Fasn PE=1 SV=2 |
| 47 | **CDC42** | Not detected | Not detected |
| 48 | **RAP1B** | Not detected | Not detected |
| 49 | **CCT2** | TCPB_HUMAN T-complex protein 1 subunit beta OS=Homo sapiens OX=9606 GN=CCT2 PE=1 SV=4 | TCPB_MOUSE T-complex protein 1 subunit beta OS=Mus musculus OX=10090 GN=Cct2 PE=1 SV=4 |
| 50 | **YWHAG** | 1433G_HUMAN 14-3-3 protein gamma OS=Homo sapiens OX=9606 GN=YWHAG PE=1 SV=2 | Not detected |
| 51 | **GNB2** | Not detected | Not detected |
| 52 | **ACTN4** | ACTN4_HUMAN Alpha-actinin-4 OS=Homo sapiens OX=9606 GN=ACTN4 PE=1 SV=2 | Not detected |
| 53 | **RAB5C** | Not detected | RAB5C_MOUSE Ras-related protein Rab-5C OS=Mus musculus OX=10090 GN=Rab5c PE=1 SV=2 |
| 54 | **C3** | Not detected | Not detected |
| 55 | **RAB10** | Not detected | Not detected |
| 56 | **HIST1H4A** | H4_HUMAN Histone H4 OS=Homo sapiens OX=9606 GN=HIST1H4A PE=1 SV=2 | Not detected |
| 57 | **KRT1** | 1) K2C1_HUMAN Keratin, type II cytoskeletal 1 OS=Homo sapiens OX=9606 GN=KRT1 PE=1 SV=6  2) K1C9_HUMAN Keratin, type I cytoskeletal 9 OS=Homo sapiens OX=9606 GN=KRT9 PE=1 SV=3  3) K1C10_HUMAN Keratin, type I cytoskeletal 10 OS=Homo sapiens OX=9606 GN=KRT10 PE=1 | E9Q1Z0_MOUSE Keratin 90 OS=Mus musculus OX=10090 GN=Krt90 PE=1 SV=1 |
| 58 | **FN1** | FINC_HUMAN Fibronectin OS=Homo sapiens OX=9606 GN=FN1 PE=1 SV=4 | Not detected |
| 59 | **AHCY** | SAHH_HUMAN Adenosylhomocysteinase OS=Homo sapiens OX=9606 GN=AHCY PE=1 SV=4 | Not detected |
| 60 | **A2M** | Not detected | Not detected |
| 61 | **BSG** | Not detected | Not detected |
| 62 | **ACTN1** | ACTN1_HUMAN Isoform 3 of Alpha-actinin-1 OS=Homo sapiens OX=9606 GN=ACTN1 | Not detected |
| 63 | **ANXA7** | Not detected | Not detected |
| 64 | **ACLY** | ACLY_HUMAN ATP-citrate synthase OS=Homo sapiens OX=9606 GN=ACLY PE=1 SV=3 | Not detected |
| 65 | **HIST1H4B** | Not detected | Not detected |
| 66 | **GDI2** | Not detected | Q3TIY6_MOUSE Rab GDP dissociation inhibitor OS=Mus musculus OX=10090 GN=Gdi2 PE=2 SV=1 |
| 67 | **FLNA** | FLNA_HUMAN Filamin-A OS=Homo sapiens OX=9606 GN=FLNA PE=1 SV=4 | Not detected |
| 68 | **UBA1** | Not detected | A0A1S6GWH5_MOUSE Uncharacterized protein OS=Mus musculus OX=10090 GN=Uba1 PE=2 SV=1 |
| 69 | **GNAS** | Not detected | Not detected |
| 70 | **GSN** | Not detected | Not detected |
| 71 | **CCT4** | TCPD_HUMAN T-complex protein 1 subunit delta OS=Homo sapiens OX=9606 GN=CCT4 PE=1 SV=4 | TCPD_MOUSE T-complex protein 1 subunit delta OS=Mus musculus OX=10090 GN=Cct4 PE=1 SV=3 |
| 72 | **RAN** | Not detected | Not detected |
| 73 | **PRDX2** | Not detected | Not detected |
| 74 | **RHOA** | Not detected | Not detected |
| 75 | **CCT3** | Not detected | TCPG_MOUSE T-complex protein 1 subunit gamma OS=Mus musculus OX=10090 GN=Cct3 PE=1 SV=1 |
| 76 | **RAC1** | Not detected | Not detected |
| 77 | **LGALS3BP** | LG3BP_HUMAN Galectin-3-binding protein OS=Homo sapiens OX=9606 GN=LGALS3BP PE=1 SV=1 | LG3BP_MOUSE Galectin-3-binding protein OS=Mus musculus OX=10090 GN=Lgals3bp PE=1 SV=1 |
| 78 | **TCP1** | Not detected | TCPA_MOUSE T-complex protein 1 subunit alpha OS=Mus musculus OX=10090 GN=Tcp1 PE=1 SV=3 |
| 79 | **KRT10** | Not detected |  |
| 80 | **CAP1** | Not detected | CAP1_MOUSE Adenylyl cyclase-associated protein 1 OS=Mus musculus OX=10090 GN=Cap1 PE=1 SV=4 |
| 81 | **RAB7A** | Not detected | Not detected |
| 82 | **TUBB4B** | 1) TBB5_HUMAN Tubulin beta chain OS=Homo sapiens OX=9606 GN=TUBB PE=1 SV=2  2) TBB4B_HUMAN Tubulin beta-4B chain OS=Homo sapiens OX=9606 GN=TUBB4B PE=1 SV=1 | Not detected |
| 83 | **HSPA5** | Not detected | BIP_MOUSE Endoplasmic reticulum chaperone BiP OS=Mus musculus OX=10090 GN=Hspa5 PE=1 SV=3 |
| 84 | **IQGAP1** | Not detected | Not detected |
| 85 | **GPI** | Not detected | Not detected |
| 86 | **RALA** | Not detected | Not detected |
| 87 | **KPNB1** | IMB1_HUMAN Importin subunit beta-1 OS=Homo sapiens OX=9606 GN=KPNB1 PE=1 SV=2 | Not detected |
| 88 | **HIST1H4I** | Not detected | Not detected |
| 89 | **TFRC** | Not detected | Not detected |
| 90 | **EIF4A1** | 1) EIF3L_HUMAN Eukaryotic translation initiation factor 3 subunit L OS=Homo sapiens OX=9606 GN=EIF3L PE=1 SV=1  2) DDX17_HUMAN Probable ATP-dependent RNA helicase DDX17 OS=Homo sapiens OX=9606 GN=DDX17 PE=1 SV=2 | 1) B7ZWF1_MOUSE Ddx3x protein OS=Mus musculus OX=10090 GN=Ddx3x PE=2 SV=1  2) IF4A3_MOUSE Eukaryotic initiation factor 4A-III OS=Mus musculus OX=10090 GN=Eif4a3 PE=1 SV=3  3) Q8VH52_MOUSE Translation initiation factor-3 subunit 5 (Fragment) OS=Mus musculus OX=10090 GN=Eif3f PE=2 SV=1 |
| 91 | **HIST4H4** | Not detected | Not detected |
| 92 | **CCT8** | TCPQ_HUMAN T-complex protein 1 subunit theta OS=Homo sapiens OX=9606 GN=CCT8 PE=1 SV=4 | Q8BVY8_MOUSE Uncharacterized protein OS=Mus musculus OX=10090 GN=Cct8 PE=2 SV=1 |
| 93 | **TLN1** | Not detected | Q80TM2_MOUSE MKIAA1027 protein (Fragment) OS=Mus musculus OX=10090 GN=Tln1 PE=2 SV=4 |
| 94 | **HIST1H4K** | Not detected | Not detected |
| 95 | **HIST1H4H** | Not detected | Not detected |
| 96 | **CCT6A** | TCPZ_HUMAN T-complex protein 1 subunit zeta OS=Homo sapiens OX=9606 GN=CCT6A PE=1 SV=3 | Not detected |
| 97 | **ANXA11** | Not detected | Not detected |
| 98 | **HIST1H4J** | Not detected | Not detected |
| 99 | **HIST1H4F** | Not detected | Not detected |
| 100 | **HIST1H4D** | Not detected | Not detected |

**Table S2**. Gene IDs for D3-EVs versus RAW-EVs

| **Gene ID** | |
| --- | --- |
| **Naïve D3-EV** | **Naïve RAW-EV** |
| GAPDH | PDCD6IP |
| ANXA2 | GAPDH |
| PKM | HSPA8 |
| HSP90AA1 | ACTB |
| ENO1 | CD9 |
| YWHAZ | PKM |
| YWHAE | ENO1 |
| CLTC | ANXA5 |
| ALDOA | HSP90AB1 |
| ALB | EEF1A1 |
| TPI1 | PGK1 |
| MYH9 | CLTC |
| SLC3A2 | PPIA |
| YWHAQ | ALDOA |
| ANXA1 | EEF2 |
| LDHA | ALB |
| FASN | TPI1 |
| CCT2 | VCP |
| YWHAG | CFL1 |
| ACTN4 | MSN |
| HIST1H4A | ATP1A1 |
| KRT1 | MYH9 |
| FN1 | SLC3A2 |
| AHCY | PFN1 |
| ACTN1 | CLIC1 |
| ACLY | ITGB1 |
| FLNA | LDHA |
| CCT4 | FASN |
| LGALS3BP | CCT2 |
| TUBB4B | RAB5C |
| KPNB1 | KRT1 |
| EIF4A1 | GDI2 |
| CCT8 | UBA1 |
| CCT6A | CCT4 |
|  | EIF4A1 |
|  | CCT8 |
|  | TLN1 |
|  | CCT3 |
|  | LGALS3BP |
|  | TCP1 |
|  | CAP1 |
|  | HSPA5 |

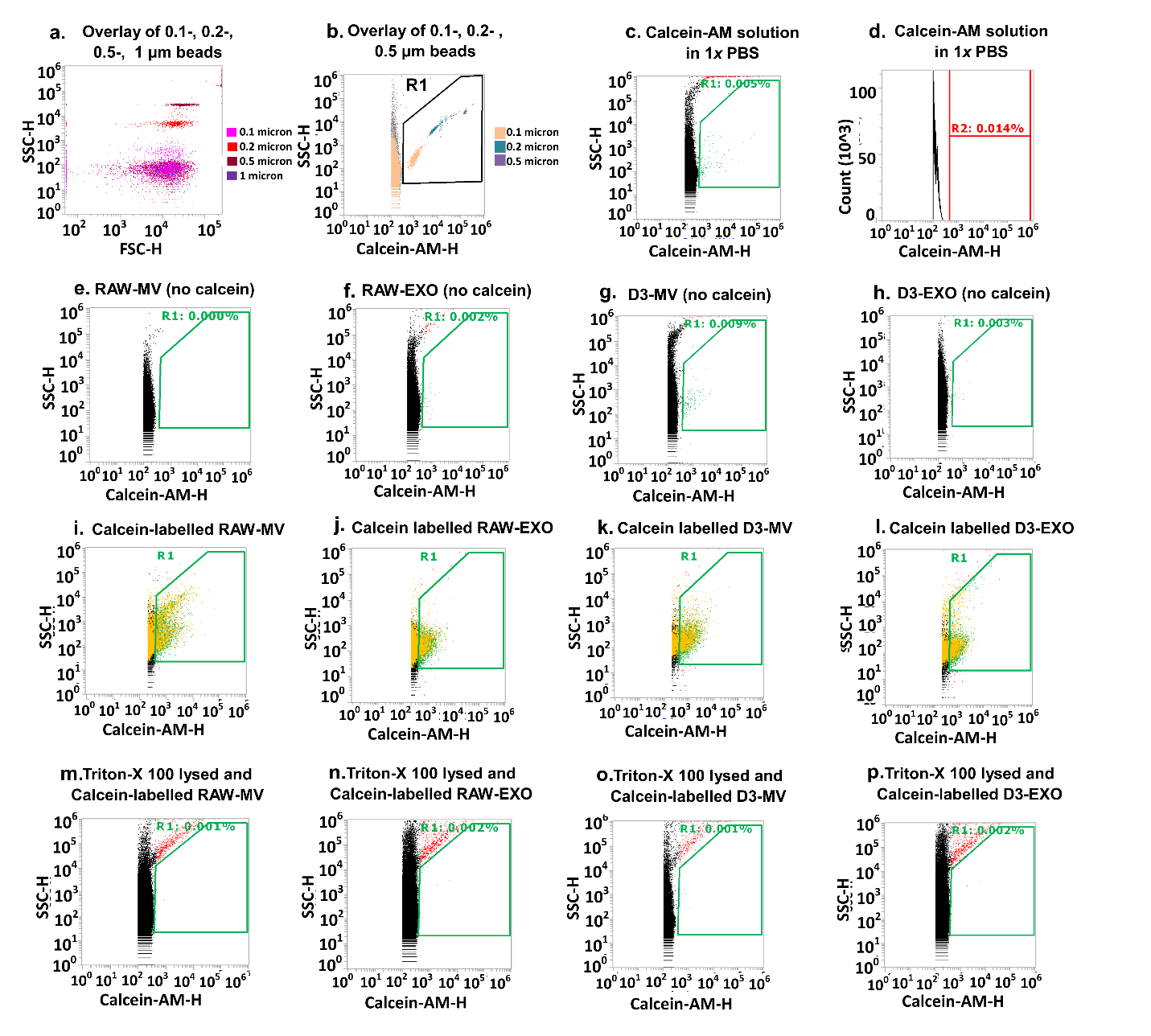

**Figure S1**. **Flow cytometry analysis of EVs**. (*a*) Side scatter (SSC)/Forward scatter (FSC) dot plots of 0.1, 0.2, and 0.5 µm microbeads obtained from an Attune NxT Acoustic Focusing Cytometer. The EV gate was defined below 0.5 µm, (*b*) SSC/Calcein-AM (BL1) fluorescent profiles of 0.1, 0.2, and 0.5 µm microbeads used to set the EV gate and establish the measurement settings, (*c*) SSC/Calcein-AM scatter plots and (*d*) histogram of fluorescent signals demonstrating separation of the instrument noise and background signals for free calcein-AM solution in 1*x* PBS. EVs were isolated from hCMEC/D3 and RAW 264.7 cell lines and resuspended in 1*x* PBS. Non-stained RAW-MV (*e*), RAW-EXO (*f*), D3-MV (*g*) and D3-EXO (*h*) were used as negative controls to verify the absence of fluorescent signals in the gated area. Scatter plots of calcein-stained RAW-MV (*i*), RAW-EXO (*j*), D3-MV (*k*) and D3-EXO (*l*). Scatter plots of calcein-stained samples treated with Triton X (1 % v/v): (*m*) RAW-MV (*n*) RAW-EXO (*o*) D3-MV and (*p*) D3-EXO. The data shown are representative plots of n=3 samples.

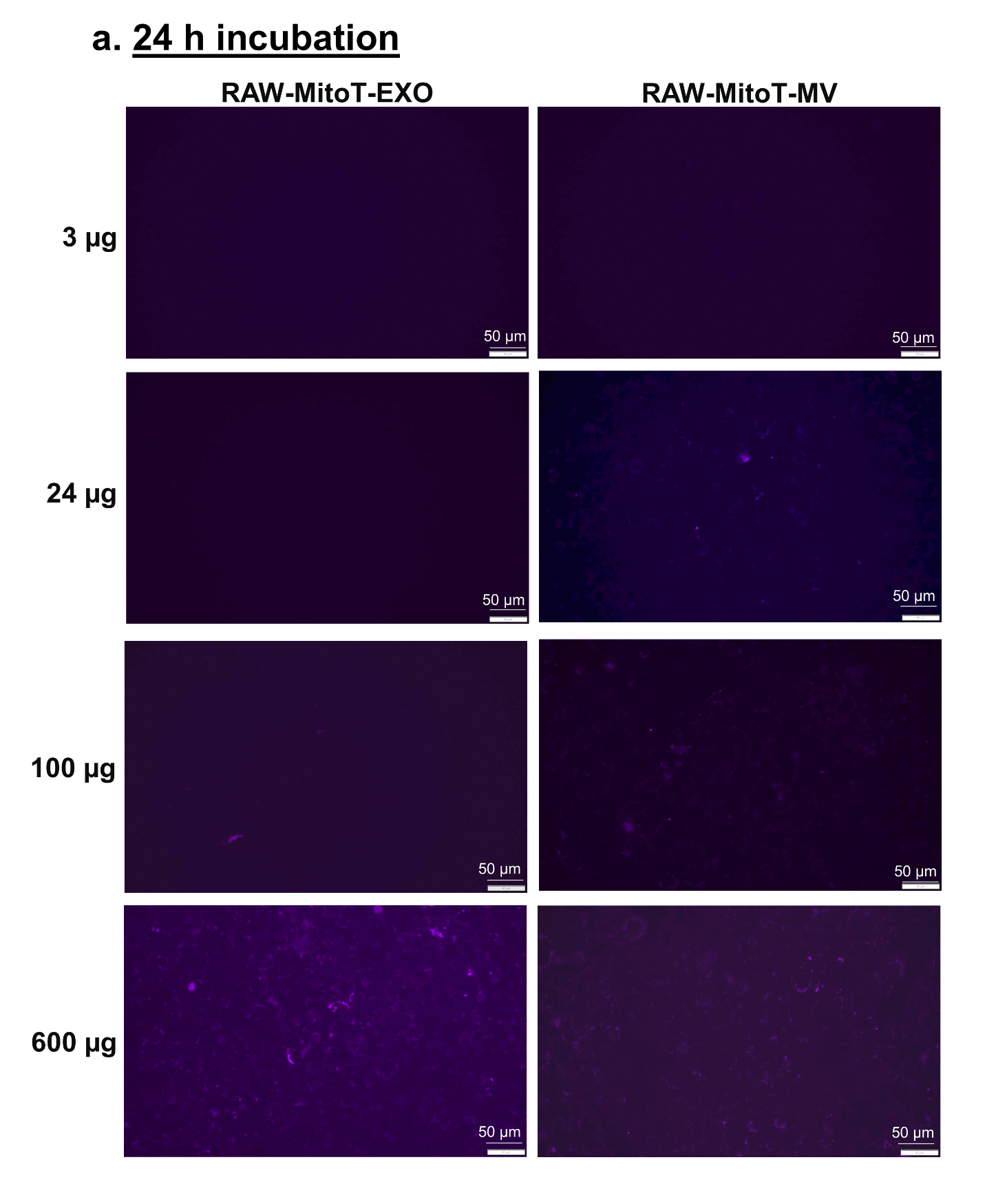

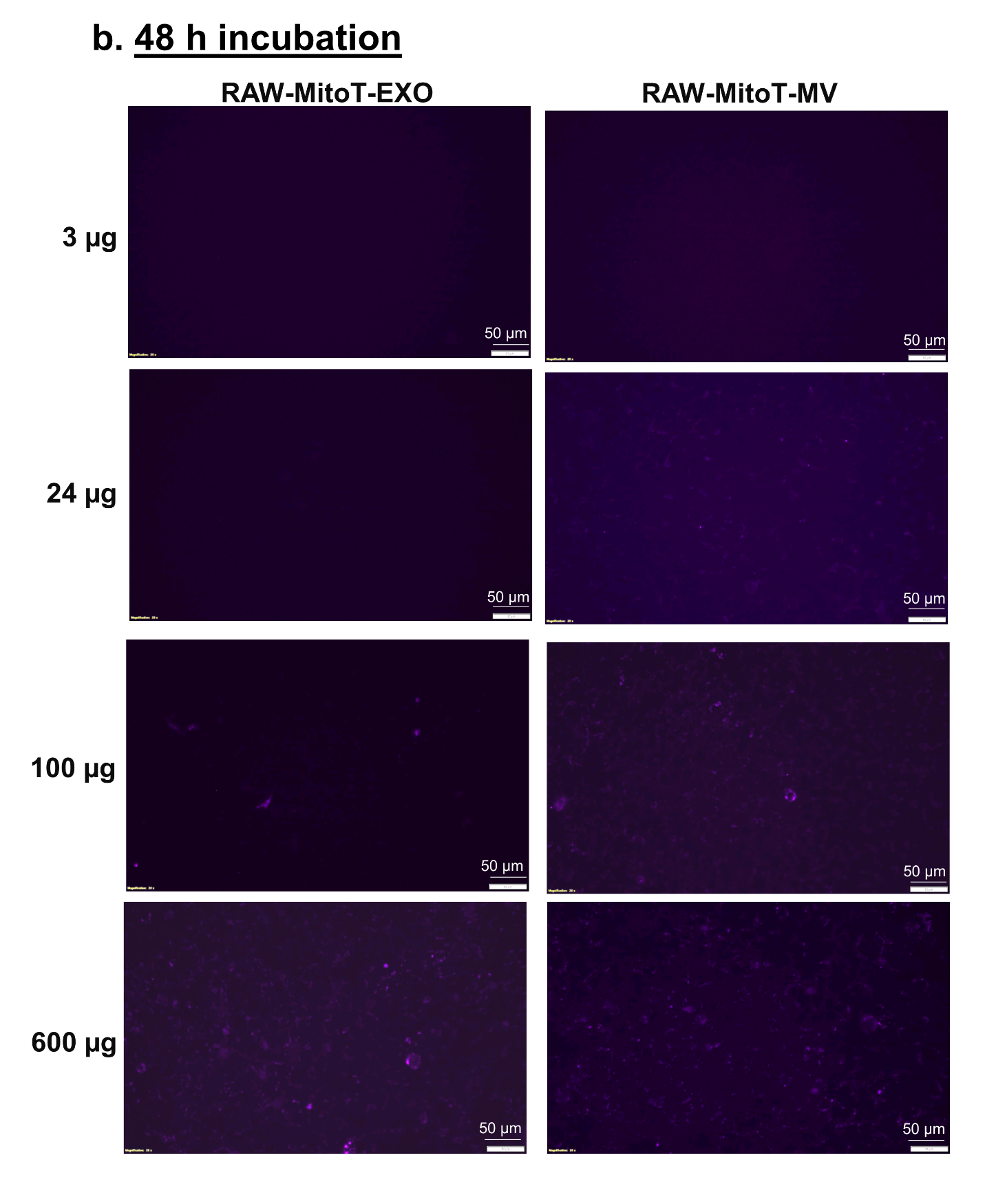

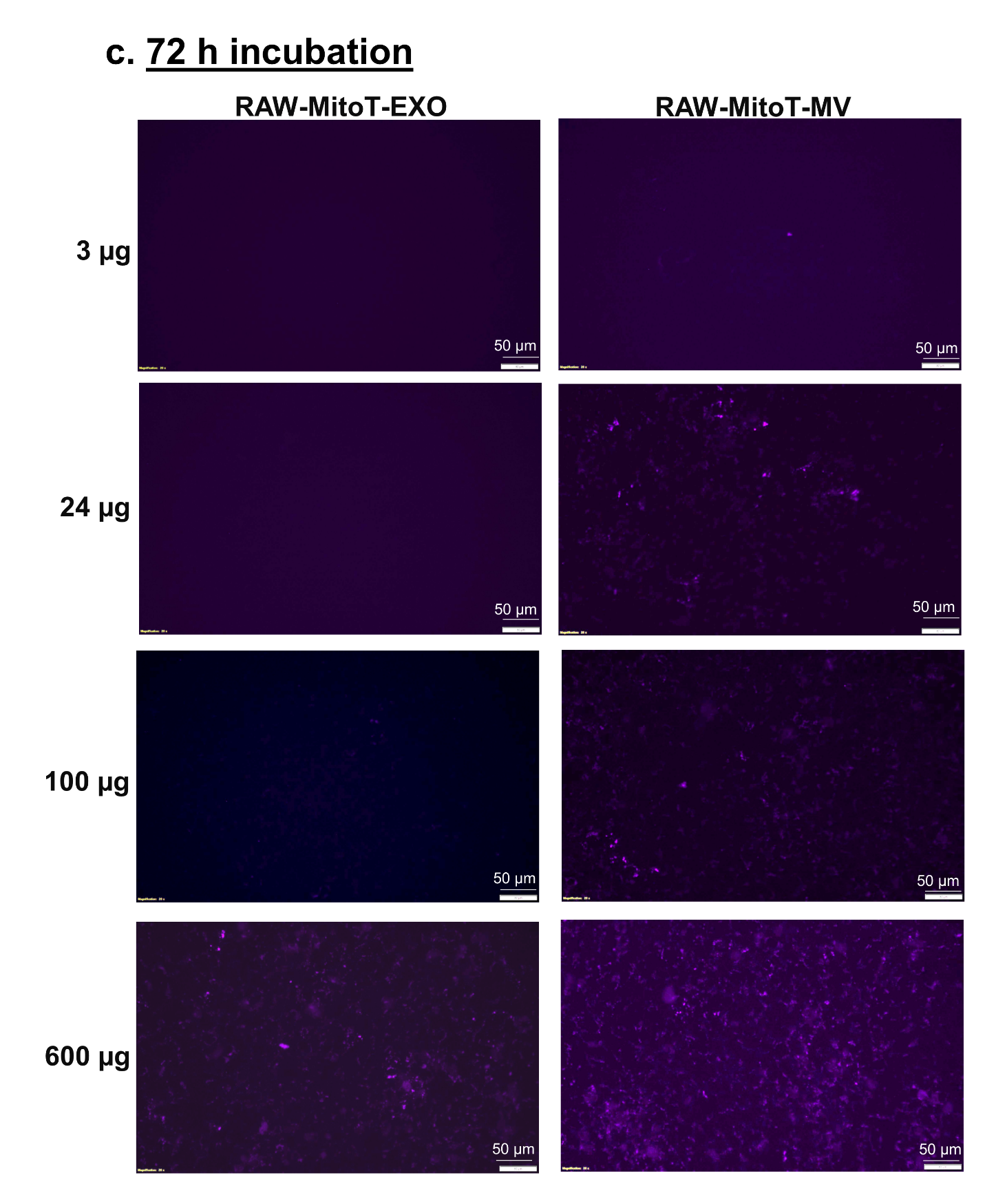

**Figure S2**. **Transfer of mitochondria from RAW-MitoT-EXO and RAW-MitoT-MV to the recipient hCMEC/D3 endothelial cells**. The donor/source RAW 264.7 macrophages were stained with MitoTracker Deep-Red (MitoT) (250 nM for 30 min) to specifically label polarized mitochondria following which the MitoT-EVs were isolated from conditioned media. The recipient hCMEC/D3 endothelial cells were treated with RAW-MitoT-EXO and RAW-MitoT-MV at the indicated protein doses and observed under an Olympus IX 73 epifluorescent inverted microscope (Olympus, Pittsburgh, PA) under the Cy5 channel settings at 24 h, 48 h and 72 h post-treatment. The presented data are representative images from three independent experiments (n=3 per experiment). Scale bar = 50 μm.

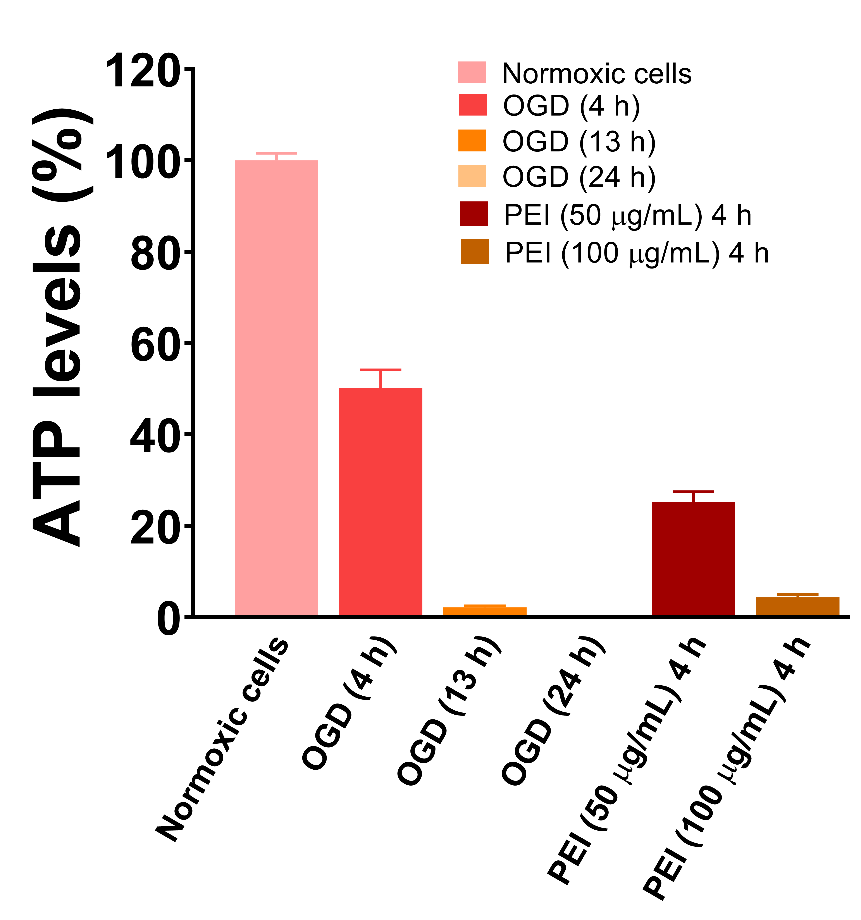

**Figure S3**. **Modelling oxygen-glucose deprivation (OGD)-induced cell death in hCMEC/D3 endothelial cells**. Confluent hCMEC/D3 endothelial cells seeded in a 96-well plate were subjected to OGD conditions for the indicated times. OGD was induced by exposing the cells in a sealed hypoxic (90% N_2_, 5% H_2_, 5% CO_2_) chamber and glucose-free media at 37 °C in a humidified incubator. An ATP assay was performed to measure the cell viability post-OGD exposure. The resulting ATP levels were compared to normoxic cells cultured in complete media. Normoxic cells treated with polyethyleneimine (PEI) for 4 h were used as a positive control. Data are represented as mean ± SD (n = 6).
